# The FGF/FGFR system in the microglial neuroinflammation with *Borrelia burgdorferi*: intersectionality with other neurological conditions

**DOI:** 10.1101/2022.08.22.504844

**Authors:** Geetha Parthasarathy, Melissa B. Pattison, Cecily C. Midkiff

**Affiliations:** Division of Immunology, Tulane University, 18703, Three Rivers Road, Covington, LA-70433, USA; Division of Microbiology, Tulane University, 18703, Three Rivers Road, Covington, LA-70433, USA; Division of Comparative Pathology, Tulane National Primate Research Center, Tulane University, 18703, Three Rivers Road, Covington, LA-70433, USA

**Keywords:** Lyme neuroborreliosis, *B. burgdorferi*, rhesus microglia, FGFR, FGF, neuroinflammation

## Abstract

**Background:** Lyme neuroborreliosis, caused by the bacterium *Borrelia burgdorferi* affects both the central and peripheral nervous systems (CNS, PNS). The CNS manifestations, especially at later stages, can mimic/cause many other neurological conditions including psychiatric disorders, dementia, and others, with a likely neuroinflammatory basis. The pathogenic mechanisms associated with Lyme neuroborreliosis, however, are not fully understood.

**Methods:** In this study, using cultures of primary rhesus microglia, we explored the roles of several fibroblast growth factor receptors (FGFRs) and fibroblast growth factors (FGFs) in neuroinflammation associated with live *B. burgdorferi* exposure. FGFR specific siRNA and inhibitors, custom antibody arrays, ELISAs, immunofluorescence and microscopy were used to comprehensively analyze the roles of these molecules in microglial neuroinflammation due to *B. burgdorferi*.

**Results:** FGFR1- 3 expressions were upregulated in microglia in response to *B. burgdorferi*. Inhibition of FGFR 1, 2 and 3 signaling using siRNA and three different inhibitors showed that FGFR signaling is proinflammatory in response to the Lyme disease bacterium. FGFR1 activation also contributed to non-viable *B. burgdorferi* mediated neuroinflammation. Analysis of the *B. burgdorferi* conditioned microglial medium by a custom antibody array showed that several FGFs are induced by the live bacterium including FGF6, FGF10 and FGF12, which in turn induce IL-6 and/or IL-8 in a dose dependent manner, indicating a proinflammatory nature. To our knowledge, this is also the first-ever described role for FGF6 and FGF12 in CNS neuroinflammation. FGF23 upregulation, in addition, was observed in response to the Lyme disease bacterium. *B. burgdorferi* exposure also downregulated many FGFs including FGF 5,7, 9, 11,13, 16, 20 and 21. Some of the upregulated FGFs have been implicated in major depressive disorder or dementia development, while the downregulated ones have been demonstrated to have protective roles in epilepsy, Parkinson’s disease, Alzheimer’s disease, spinal cord injury, blood-brain barrier stability, and others.

**Conclusions:** In this study we show that FGFRs and FGFs are novel mediators of inflammatory pathogenesis in Lyme neuroborreliosis. It is likely that an unresolved, long-term (neuro)-Lyme infection can contribute to the development of other neurologic conditions in susceptible individuals either by augmenting pathogenic FGFs or by suppressing ameliorative FGFs or both.

## INTRODUCTION

Tick-borne infections account for 77 - 95% of all vector-borne diseases in the United States. Of these, Lyme disease (LD) is the leading tick-borne illness in the northern hemisphere accounting for 70% of all reported tick-borne diseases [1]. Caused by the gram-negative bacterium *Borrelia burgdorferi*, the annual case load of LD is ~476000 cases [2], up from the previous estimates of 300,000 per year [3]. Lyme neuroborreliosis (LNB) is a form of Lyme disease that affects both the central and peripheral nervous systems (CNS, PNS), and accounts for ~15-25% of all the LD cases. Signs and symptoms of LNB range from meningitis, cranial neuritis, radiculoneuropathies, encephalitis, vasculitis (rarely) in the early stages, to a broad range of neuropsychiatric/neuropsychological conditions including anxiety, depression, cognitive impairment, obsessive compulsive disorders, schizophrenia and dementia-like syndromes in the later stages [4]. While depression is a common late stage manifestation (22%-60% of LNB cases [4]), dementias are rare and make up to 6% of LNB sequelae [5]. Interestingly, other than secondary dementias associated with LNB, presence of the organism or Lyme infection has also been documented in patients with Alzheimer’s disease (AD)-like pathology, Parkinson’s disease (PD), Lewy Body dementia (LBD) and fronto temporal dementia (FTD) [6–10]. Whether this association is correlation or causation has been a matter of debate. It is possible that commonalities in pathogenesis exist between LNB and these diseases, and these commonalities can cause Lyme infection to augment/contribute towards other neurological diseases, or result in disease-like pathologies. However, identification of such commonalities requires understanding the pathogenesis of diseases in question and decipher the intersectionality.

In recent years, the FGFR/ FGF system has been widely studied in several neurological diseases including AD, PD, depression, anxiety, multiple sclerosis, epilepsy, schizophrenia and others [11–17]. The FGFR family comprises 4 receptors FGFR1-4, which are transmembrane tyrosine kinases. Their ligands are FGFs, 22 in number, of which 18 are known to bind FGFRs. Signaling via FGFR is thought to be neuroprotective and to dampen neuroinflammation [18]. For this reason, FGFR agonists have been considered as therapeutic targets in AD, PD, traumatic brain injury and others [19]. However, neurotoxic effects have also been observed, with FGFR signaling mediating apoptosis in amyotrophic lateral sclerosis (ALS) [20], and axon degeneration in experimental autoimmune encephalitis (EAE), [17] indicating divergent roles in different neurological diseases.

Since many of the conditions/symptoms studied with respect to FGFR overlap with LNB and its sequelae, FGFR system as a possible commonality between Lyme infection and other neurological conditions seemed intriguing. Therefore, we decided to investigate the role of FGF/FGFR system in primary rhesus microglia, the most significant mediator of neuroinflammation in the CNS. Since microglia only comprise ~6-10% of the total glial cells, they are rare [21]. Un-inoculated, young rhesus brain tissues from which microglia are extracted and cultured are just as rare. By using these scarce resources, siRNA, several inhibitors, custom antibody arrays, immunofluorescence and immunoassays we have built a detailed picture of the FGF/FGFR system in microglial neuroinflammation due to *B. burgdorferi*. <u>To our knowledge, this is the first **comprehensive** FGF/FGFR study, both for Lyme disease and bacteria in general.</u> In a study that took over four years to complete, we provide a valuable insight into how a neurological bacterial infection can contribute or exacerbate other neurological diseases/conditions and likely affect treatment modalities.

## MATERIALS AND METHODS

### Bacterial strain and culture

*B. burgdorferi* strain B31, clone 5A19, was cultured according to previously published protocols [22]. Briefly, bacteria were cultured under microaerophilic conditions in Barbour-Stoenner-Kelly (BSK-H) medium supplemented with amphotericin (0.25 μg/mL), phosphomycin (193 μg/mL) and rifampicin (45.4 μg/mL), for about 5-6 days. (All from Millipore Sigma, St. Louis, MO). A dark field microscope was used to determine bacterial concentration, and required number of bacteria was harvested by centrifugation at 2095 x g for 30 minutes at room temperature (without brakes). The bacterial pellet was resuspended in DMEM: F12 (ThermoFisher Scientific, Waltham, MA) supplemented with 10% fetal bovine serum (FBS, Hyclone, Fisher Scientific, Hampton, NH) to the same concentration prior to pelleting. For the experiments, bacteria were diluted further in the same medium supplemented with 0.5 ng/mL granulocyte macrophage colony stimulating factor (GM-CSF, Millipore Sigma), to the required multiplicity of infection (MOI).

### Isolation and culture of primary microglia

Primary microglia were isolated from frontal cortex tissues of rhesus macaques (*Macaca mulatta*) as described previously [22]. Briefly, brain tissues were obtained from un-inoculated young animals from the breeding colony that were euthanized due to injury or persistent idiopathic diarrhea. <u>Euthanasia protocols, all performed by veterinarians, were approved by the Tulane Institutional Animal Care and Use Committee (Tulane IACUC).</u> The leptomeningeal blood vessels and the leptomeninges were removed first with fine tweezers, followed by mincing of the tissue with scalpels. The finely minced tissue was then subjected to enzymatic digestion with 0.25% Trypsin-EDTA containing 200 Kunitz unit/mL DNaseI (Sigma Aldrich, St. Louis-MO) at 37°C for 20 minutes. Following digestion, the tissue was centrifuged at 335 x g, for 10 minutes, upper layer of cells removed and filtered through a 20μm Nitex filter. The filtrate was resuspended in DMEM: F12 supplemented with 10% FBS, 1% penicillin-streptomycin and 0.5 ng/mL GM-CSF. The aggregate cultures were seeded in T-75 flasks and incubated at 37°C, 5% CO_2_. Medium was changed every four days for about 4 weeks, prior to harvesting of microglia. Microglia were isolated by vigorous tapping of the sides of the T-75 flasks, counted and seeded at the desired density. Typical yield of microglia was between 90-95%, unless otherwise stated. Microglial identity was verified by microglial marker Iba1 (1:10 to 1:25-mouse monoclonal #sc-32725, Santa Cruz Biotechnology; 1:100-rabbit polyclonal, #019-19741, FujiFilm Wako Pure Chemical Corp., Richmond, VA), as well as relative cellular size. All cell assays were conducted 2-3 days after seeding. Microglia were isolated from 9 frontal cortex tissues, obtained from animals ranging in age from 1.21 to 6.26, through the multi-year course of this study.

### RNAi

Silencing of the FGFR transcripts by siRNA was carried out as follows. Microglia were seeded on 24-well plates at a density of ~2 × 10^4^/ well. Cells were allowed to adhere for 48h (37°C, 5% CO_2_), after which medium was removed and replaced with 100 μL antibiotic-free medium. siRNA-transfection reagent complexes were generated using 2 μL HiPerfect transfection reagent (Qiagen, Germantown, MD) and 25-50 nM siRNA (non-specific control siRNA (sc-37007) or FGFR specific (FGFR1/Flg-sc-29316, FGFR2/Bek-sc-29218, FGFR3-sc-29314); Santa Cruz Biotechnology) in antibiotic and serum-free medium. The complexes were allowed to incubate at room temperature for 30 minutes, and 100 μL of the complex was added to each well. Cells containing the transfection complexes were incubated at 37°C, 5% CO_2_ for 6h, followed by addition of 400 μL of antibiotic-free medium. After a further 18h incubation, *B. burgdorferi* (MOI 10:1) or medium alone was added. Cells were incubated for an additional 24h, prior to collection of supernatants (3000 rpm, 10 minutes at 4°C).

### Infection assays with FGFR inhibitors

Microglia were seeded on 24-well plates or 4-well chamber slides at a density of ~2 x10^4^ cells/well. After 48h, cells were pretreated with specific FGFR inhibitors or solvent control (dimethyl sulfoxide (DMSO)) for about 2 h. The medium was discarded and fresh medium without antibiotics containing *B. burgdorferi* at an MOI of 10:1 was added, followed by addition of inhibitors or DMSO. Medium only group served as controls. After 24h at 37°C, 5% CO_2_, supernatants were collected as before and stored at −20°C until analysis. The following inhibitors were used-FGFR1 inhibitor PD166866 (#341608-Millipore Sigma); FGFR1-3 (and likely FGFR4) inhibitor BGJ398 (#HY13311-MedChem Express, Monmouth Junction, NJ); FGFR1-3 inhibitor (and likely FGFR4) AZD4547 (#HY13330-MedChem Express).

To determine whether secreted factors trigger FGFR activation, supernatants after infection assays were collected as before. They were thawed, re-centrifuged, and filtered through a 0.22 μm filter and applied to freshly cultured microglia from the same tissue grown on chamber slides. Cells were fixed after 24 h for immunofluorescence. Microglial-conditioned medium without the bacteria was similarly collected and used as a negative control.

To determine the effect of FGFs on inflammatory mediator production, various doses of specific FGFs were added to fresh microglia for 24h and supernatants, and cells analyzed as before. PBS/BSA (0.1%) was used as a solvent control. Recombinant human FGFs (FGF6 #238F6-025; FGF10-#345-FG-025; FGF12-2246-FG-025) were purchased from R&D systems (Minneapolis, MN).

### Immunofluorescence (IF)

IF was carried out as described previously [23] on experiments carried out in chamber slides. At the end of the experimentation period, supernatants were removed, and cells were fixed in ice-cold 2% paraformaldehyde for 10 minutes at room temperature on a shaking platform. Cells were briefly washed three times in cold PBS, followed by permeabilization in ethanol:acetic acid mixture (2:1) at 4°C for 5 minutes. Cells were washed again as before and kept in the same medium at 4°C until analysis with specific antibodies.

For the immunostaining, cells were re-permeabilized in PBS containing 0.1% Triton-X-100 for 15 minutes at room temperature on a shaking platform. The slides were then blocked with PBS containing 10% normal goat serum (NGS) (NGS buffer) for 1 h, followed by staining with specific primary antibody for another hour. Cells were then probed with an appropriate secondary antibody conjugated to Alexa 488 (green) or Alexa 568 (red) (1:1000, Invitrogen) for 1 h, to visualize the target protein of interest. Nuclear staining was carried out with DAPI (5 minutes, 1:5000, Millipore Sigma) as required. All the antibodies were suspended in the NGS buffer with incubations at room temperature. The following anti-human primary antibodies were used. Anti-FGFR1 (sc-121), anti-FGFR2 (sc-122), anti-FGFR3 (sc-123) (1:50; all rabbit polyclonal, Santa Cruz biotechnology) anti-phosphoFGFR1 (Tyr 653,654) (1:50; rabbit polyclonal; #44-1140G- ThermoFisher Scientific), anti-FGF6, anti-FGF10, anti-FGF12 and anti-FGF23 (all 1:50; rabbit polyclonal; FGF6-#MBS2007292, FGF10-#MBS9606991, FGF12-#MBS2028698, FGF23-#MBS9605052, MyBiosource, San Diego, CA). Slides were mounted with an anti-quenching medium, covered with cover slips and visualized for microscopy.

### Antibody-array

A custom antibody array for specific FGFs was carried out to identify the likely FGFs induced by *B. burgdorferi* exposure. Assay was conducted with RayBiotech custom L-series human array (RayBiotech, Peachtree corner, GA). The assay uses a semi-quantitative modified ELISA procedure wherein the proteins in the sample are directly labelled with biotin and used as a probe to bind corresponding antibodies printed on a glass slide. Biotin-labelled bound proteins are identified using streptavidin conjugated to fluor, and read using a laser scanner (Axon GenePix), where approximate Units of expression can be obtained. The normalized Units were then used to create semi-quantitative proteomic charts using Microsoft Excel^®^. To generate a Heatmap, the biomarker values were standardized (centering and scaling) by subtracting the average and then dividing by the standard deviation. The standardized data were plotted in a heatmap with hierarchical clustering by Euclidean distance, using the R programming language V3.6.3 (R Core Team 2017) software.

### Microscopy

FGFR, pFGFR1 and specific FGF expression in microglia were visualized using a Leica DMRE fluorescent microscope (Leica microsystems, Buffalo Grove-IL) and Lumecor SOLA GUI software (Lumencor, Beaverton-OR). Cells were imaged using the Nuance Multispectral Imaging System (CRi, PerkinElmer, Waltham-MA). Percentage of specific FGFR positive cells were counted over 5-10 frames each and graphed using Microsoft Excel^®^. Confocal microscopy was carried out using a Leica TCS SP8 confocal microscope, equipped with four lasers: 405nm (UV), argon-krypton 488 nm (blue), DPSS 561 nm (yellow), helium-neon 633 nm (far red). Adobe^®^ Photoshop CS6 was used to assemble the images.

### Quantitation of chemokines and cytokines

Custom Procartaplex-multiplex kits (ThermoFisher Scientific) were used to analyze the levels of IL-6, IL-8 and MCP-1 in samples. Assays were carried out according to manufacturer’s instructions, using Bio-Plex® 200 Suspension Array System and Bio-Plex® Manager Software Version 6.2 (Bio-Rad Laboratories, Hercules, CA). FGF6, FGF12 enzyme linked immunosorbent assays (ELISA) were carried out using calorimetric human ELISA kits (MBS454039, MBS8802366, MyBiosource). The results were graphed using Microsoft Excel^®^ and figures were assembled using Microsoft Powerpoint^®^ and Adobe^®^ Photoshop CS6.

### Statistics

For all experiments excluding the antibody array, a Student’s t-test (2-tailed) was used to determine the statistical significance of an outcome. All analyses were carried out in duplicate. A value of p < 0.05 was considered statistically significant. For the antibody array, a principal component analysis was carried out using RayBiotech statistical services.

## RESULTS

### Exposure to *B. burgdoferi* upregulates FGFRs and associated signaling pathways in primary rhesus microglia

Primary rhesus microglia were exposed to live *B. burgdorferi* for 24h and analyzed for FGFR1, 2, and 3 expressions by immunofluorescence. The results are shown in Fig. 1. Fig.1a shows the percent of microglia derived from three different animal tissues, respectively, that express specific FGFRs. Expression levels varied among tissues, but were significantly higher than medium alone controls across all tissues. Immunofluorescence photographs of FGFR1, FGFR2 and FGFR3 expression in microglia in response to *B. burgdorferi* is shown in Fig.1b, and the relative increase in expression over medium controls for FGFR2 and FGFR3 is shown in Fig.1c. Fig.1c also shows that FGFR2 and 3 expressions (green) is confined to microglia. This is shown through Iba1 staining (red). Other than Iba1 as a marker for microglial specificity, confirmation was also through the relative size of these cells. Microglia are the smallest of the glial cells and can generally be distinguished by their relatively small size, as shown in online supplementary material 1 (SM1).

**Fig. 1.**
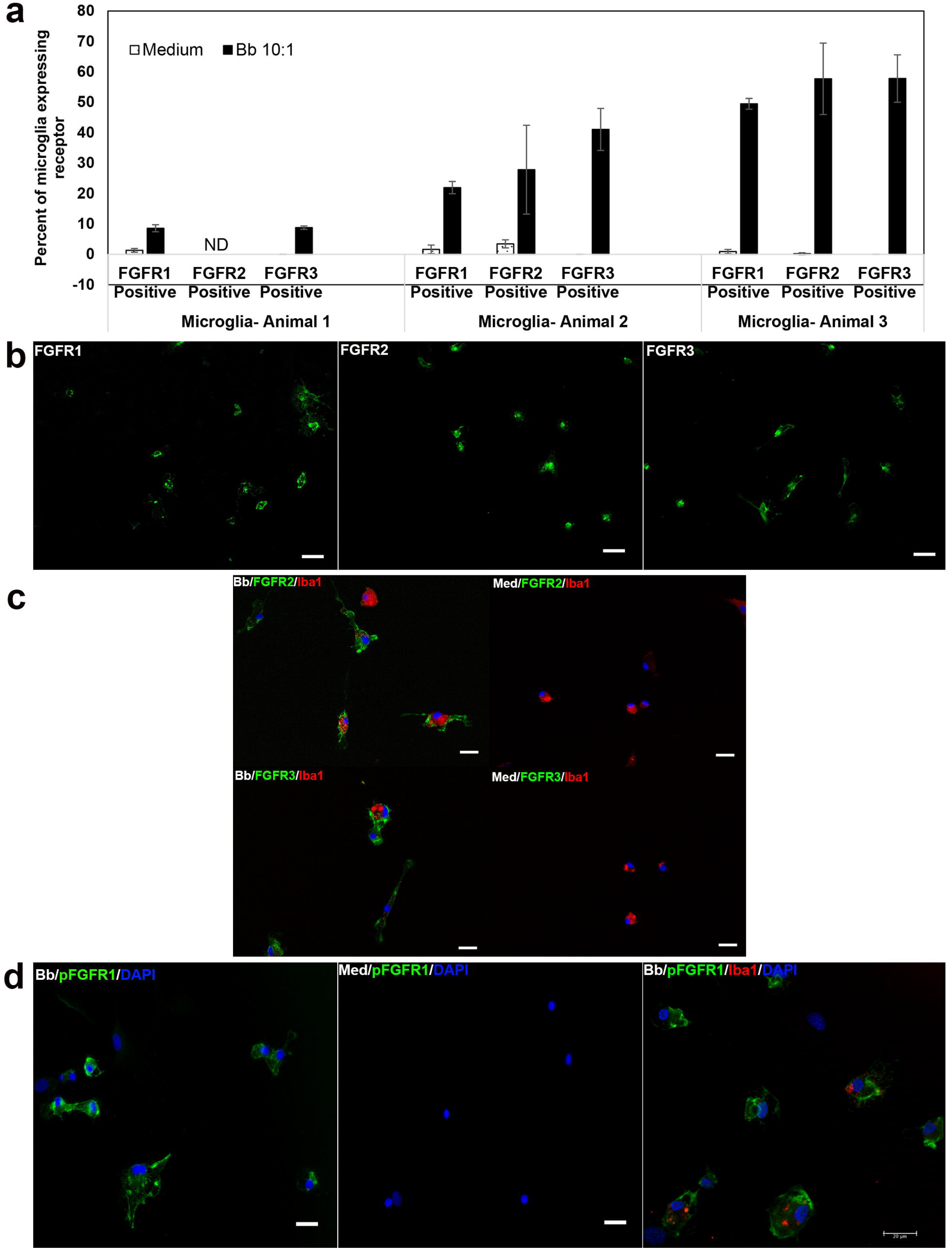
Expression of FGFRs in primary rhesus microglia in response to live *B. burgdorferi* exposure. **a**) Primary microglial cells were exposed to *B. burgdorferi* for 24h. Cells were fixed as described in Methods and stained for FGFR1, FGFR2 or FGFR3 by immunofluorescence. Percent of microglial cells expressing receptors from tissues of 3 different animals were semi-quantitated and graphed. Bar represents standard deviation. ND-not determined. **b**) Immunofluorescence microscopy pictures of FGFR expression in microglia exposed to *B. burgdorferi*. FGFR1 is from animal 1, while FGFR2 and FGFR3 expressions are from Animal 3 in **a**. Bar represents 50 μm. **c**) Immunofluorescent pictures of FGFR2 and FGFR3 staining (green) confirming the expression to be in microglia by additional staining for Iba1 (red). Microglia were derived from a fourth animal tissue. Nuclei are stained blue with DAPI. Increased expression over medium control is also seen. Bar represents 25 μm. **d**) Activation of the FGFR1 pathway is shown by increased expression of phosphoFGFR1 (pFGFR1, green) in microglial cells exposed to *B. burgdorferi* over medium controls. Bar represents 25 μm. The panel on the far-right shows confocal micrograph of microglia dually stained for Iba1 (red) and pFGFR1 (green) along with the nuclear stain DAPI in blue.

Since the expression of receptors was upregulated in response to infection, we next sought to confirm if downstream signaling is also activated. FGFRs are receptor tyrosine (Tyr) kinases that get phosphorylated at the intracellular tyrosine kinase domains, thus resulting in cell signaling. While there are several autophosphorylation sites, Tyr residues 653 and 654 are considered important for cell signaling and biological responses [24]. Therefore, phosphoFGFR1 (pFGFR1) at Tyr 653, 654 domains were also measured by immunostaining. Fig. 1d shows increased pFGFR1 in primary rhesus microglia upon exposure to live *B. burgdorferi* indicating pathway activation. While the antibody is specific for FGFR1 phosphorylation, it is to be noted that the Tyr653,654 domains are conserved across all FGFR1-4 receptors [24].

### FGFR pathways are proinflammatory in rhesus microglia in response to the Lyme disease bacterium (or its sonicated components)

To determine the effect of FGFR activation that occurred in response to the Lyme disease bacterium, RNA interference by means of siRNA was initially used. Fig 2a shows that inhibition of individual FGFR1, 2 or 3 receptors in the presence of bacteria down regulate the expression of IL-6, IL-8 and MCP-1 at 50 nM siRNA concentration. Even at the lower siRNA concentration of 25 nM, inhibition of FGFR1, 2 or 3 significantly downregulated both IL-6 and MCP-1, while only FGFR3 inhibition at this concentration affected IL-8, indicating dose dependent effects on specific mediators (not shown). SiRNA (50 nM) when used with medium alone, did not have an appreciable effect on IL-6 (or IL-8 levels for the most part), while it did have an effect on MCP-1 expression, indicating that this mediator is continuously induced at a low level in the absence of any stimuli through these receptors. To confirm the proinflammatory effect of FGFR activation in response to *B. burgdorferi*, three other FGFR inhibitors were also used and are shown in Fg.2b and Fig. 3. PD166866 is considered as an FGFR1 inhibitor, while both BGJ398 and AZD4547 are considered to be potent inhibitors of FGFRs 1-3, although they might affect FGFR4 weakly. All the inhibitors affect tyrosine kinase activity, hence autophosphorylation and signaling [25–27], Treatment of microglial cells with FGFR inhibitors showed that they had efficacies at different doses. PD166866/FGFR1 inhibitor, in the presence of *B. burgdorferi*, did not have an appreciable effect at 500 nM concentration while at higher concentrations (≥ 1μM) it significantly downregulated IL-6, IL-8 and MCP-1 (Fig. 2b). FGFR1-3 inhibitor BGJ398 on the other hand was very effective in downregulating all three mediators at 500 nM, (Fig. 3a) while the other FGFR1-3 inhibitor AZD4547 was only effective in significantly suppressing all three mediators at 5 μM and higher (Fig. 3b). This indicates that range of inhibition (FGFR1 only vs all 3) and formulation differences (affecting same targets) likely mediate the potency of the inhibitors. Only the non-toxic doses are shown. Toxicity was determined separately through an MTT based cell viability assay (not shown).

**Fig. 2.**
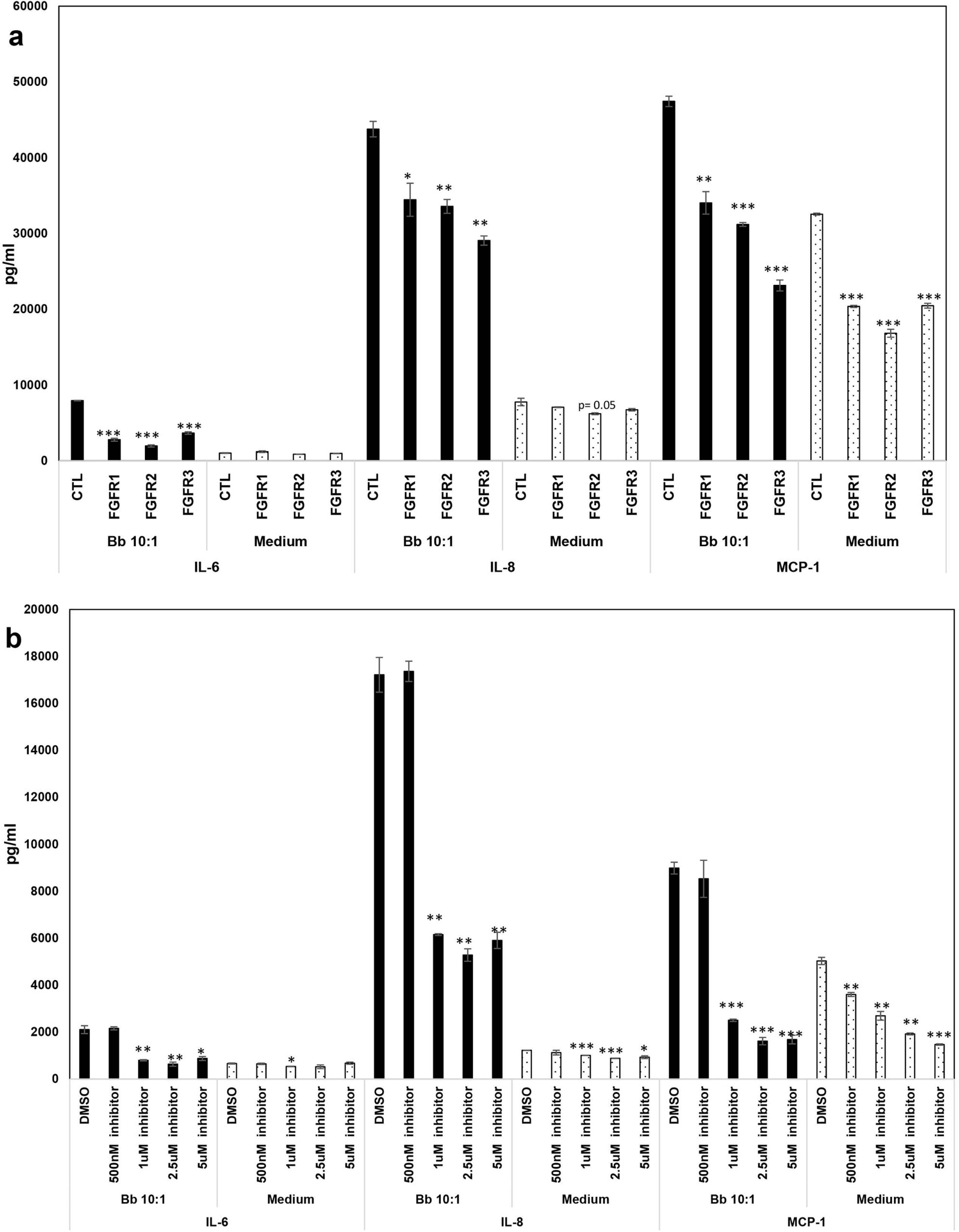
Effect of FGFR specific siRNA and FGFR1 inhibitor PD166866 on chemokine and cytokine expression by primary rhesus microglia. The effect of 50nM siRNA (control siRNA or FGFR specific siRNA) on the secretion of IL-6, IL-8 and MCP-1 is shown in (**a)**. Three experiments from microglia derived from tissues of 3 different animals were conducted. A representative graph for siRNA effect on *B. burgdorferi* induced inflammatory mediator secretion is shown. siRNA effect on medium controls from the same tissue is included. **b**) shows the effect of FGFR1 inhibitor PD166866 on inflammatory mediator output from primary rhesus microglia in response to *B. burgdorferi*. A representative experiment is shown for *B. burgdorferi* along with medium controls from the same animal tissue. Three experiments were carried out on microglia derived from tissues of two different animals. Bar represents standard deviation for both **a** and **b**. All statistical comparisons are with Control siRNA or DMSO within each treatment group (*B. burgdorferi* or Medium). * p< 0.05, **p < 0.01, and *** p< 0.001.

**Fig. 3.**
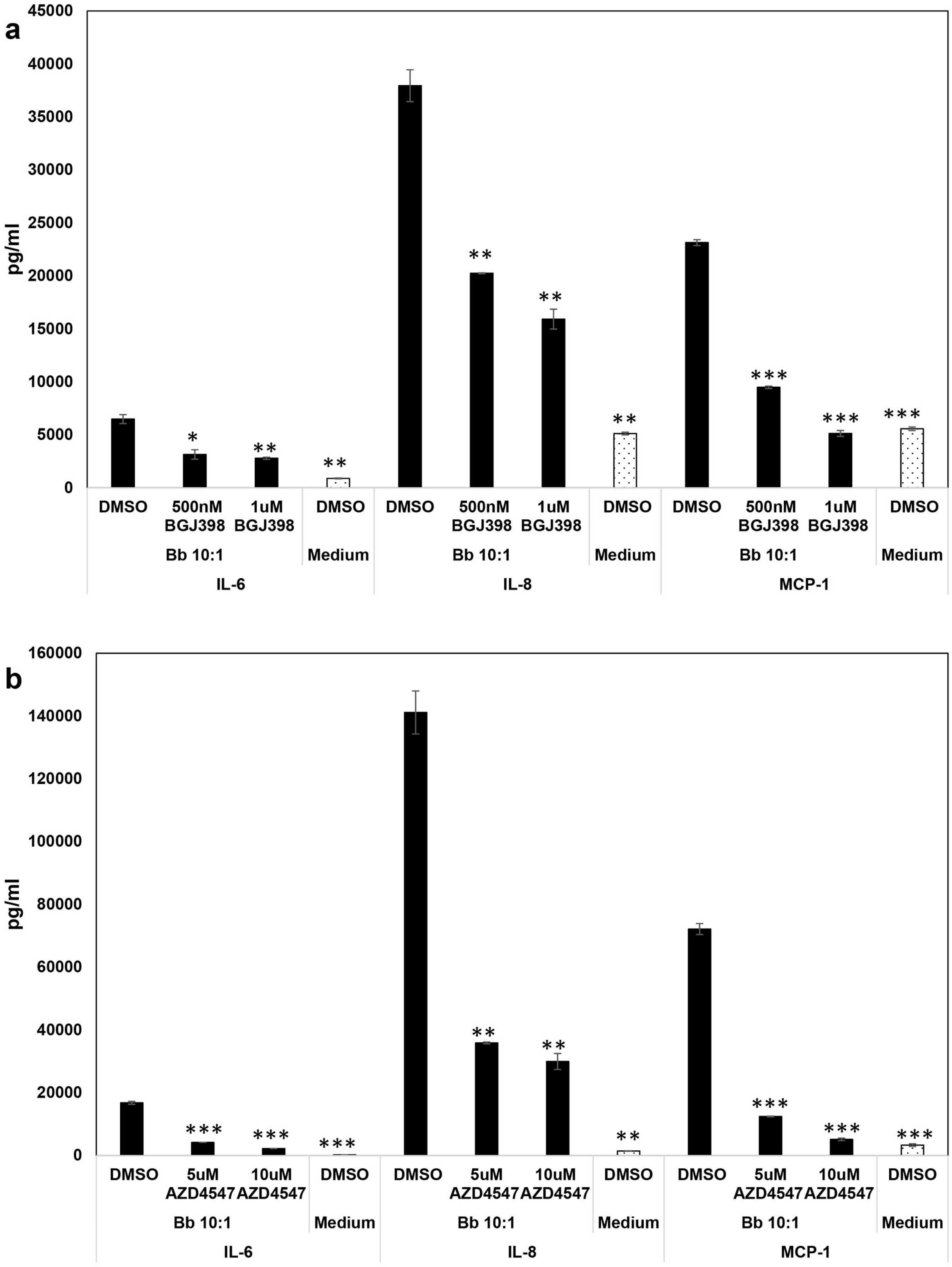
Effect of FGFR1-3 inhibitors BGJ398 and AZD4547 on chemokine and cytokine expression from primary rhesus microglia. Primary rhesus microglia were pretreated with various concentrations of BGJ398 (**a**) or AZD4547 (**b**) for 1 and a half to two hours prior to treatment with *B. burgdorferi*. DMSO was included as the solvent control for the drug treatments. After 24 h, supernatants were collected and analyzed for IL-6, IL-8 and MCP-1 by multiplex ELISA. Three-four experiments on microglia derived from 3 tissues were conducted for BGJ398, while 3 experiments from microglia derived from 3 tissues were carried out with AZD4547. One of the experiments is shown for each. Bar represents standard deviation. All statistical comparisons are with *B. burgdorferi* + DMSO. * p< 0.05, **p < 0.01, and *** p< 0.001.

To ensure the efficacy of the inhibitors in downregulating signaling in microglia, pFGFR1 immunostaining was conducted as before and showed that the inhibitors were effective in downregulatng the same (Supplemental material 2 (SM2)). A representative experiment for each inhibitor in the presence of *B. burgdorferi* is shown in Figs. 2 and 3, and the overall effect across all experiments is shown in Table 1. The average fold-downregulation of IL-6, IL-8 and MCP-1 in the presence of various inhibitors and siRNA across multiple experiments and multiple tissues shows that FGFRs are potent mediators of neuroinflammation in primary microglia and could be important novel pathogenic determinants in Lyme neuroborreliosis. Even without the presence of *B. burgdorferi*, they seem to mediate MCP-1 induction at a low level as seen with its significant downregulation in medium alone controls with all the inhibitors (Table 1 and supplementary material 3a (SM3A)). While the table shows the fold down-regulation in inflammation in *B. burgdorferi* and medium controls with inhibitors, Figs. 2 and 3 show that the magnitude of induction is significantly different between the two treatments and that a similar fold-downregulation with either treatment does not translate the same.

**Table 1:**
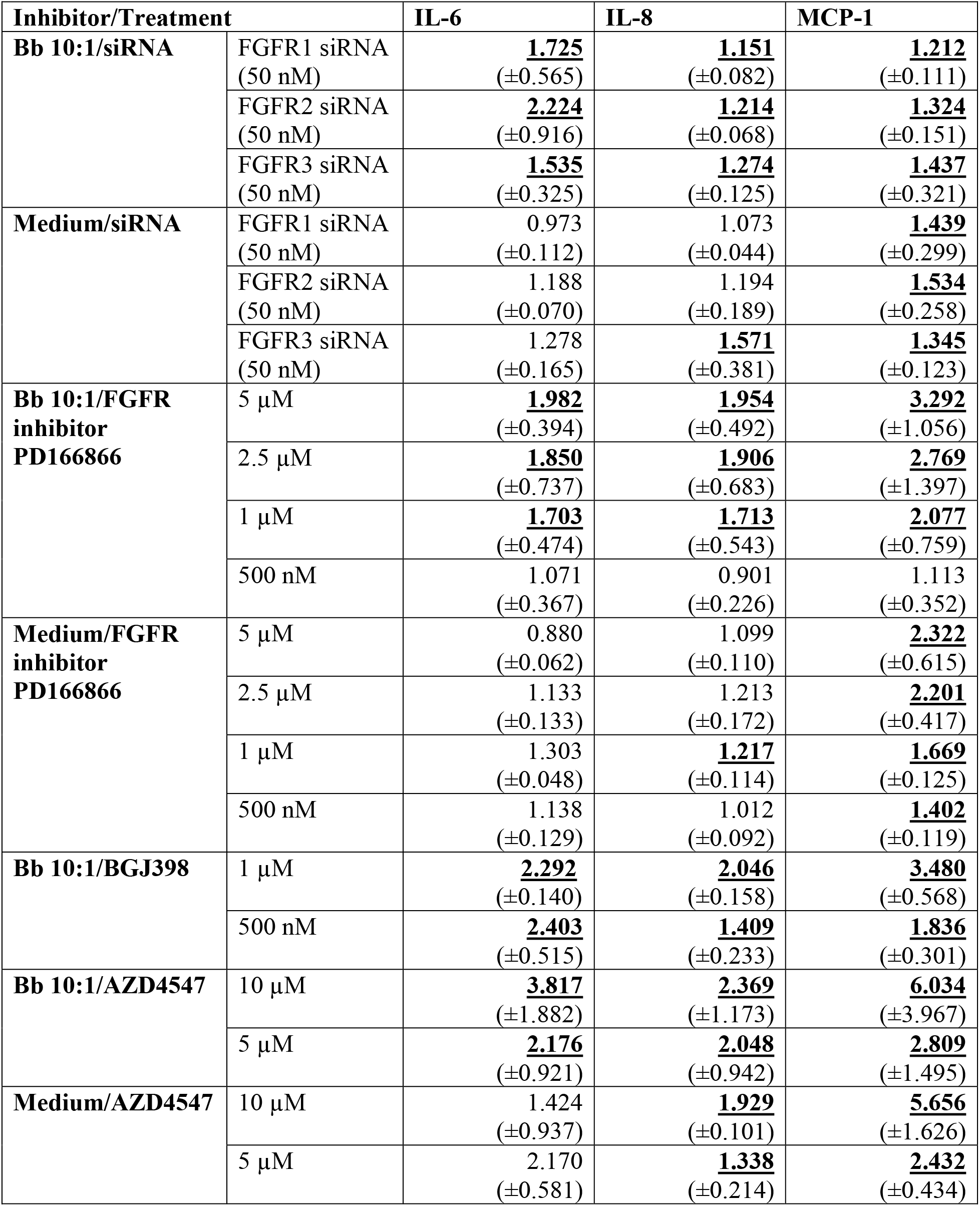
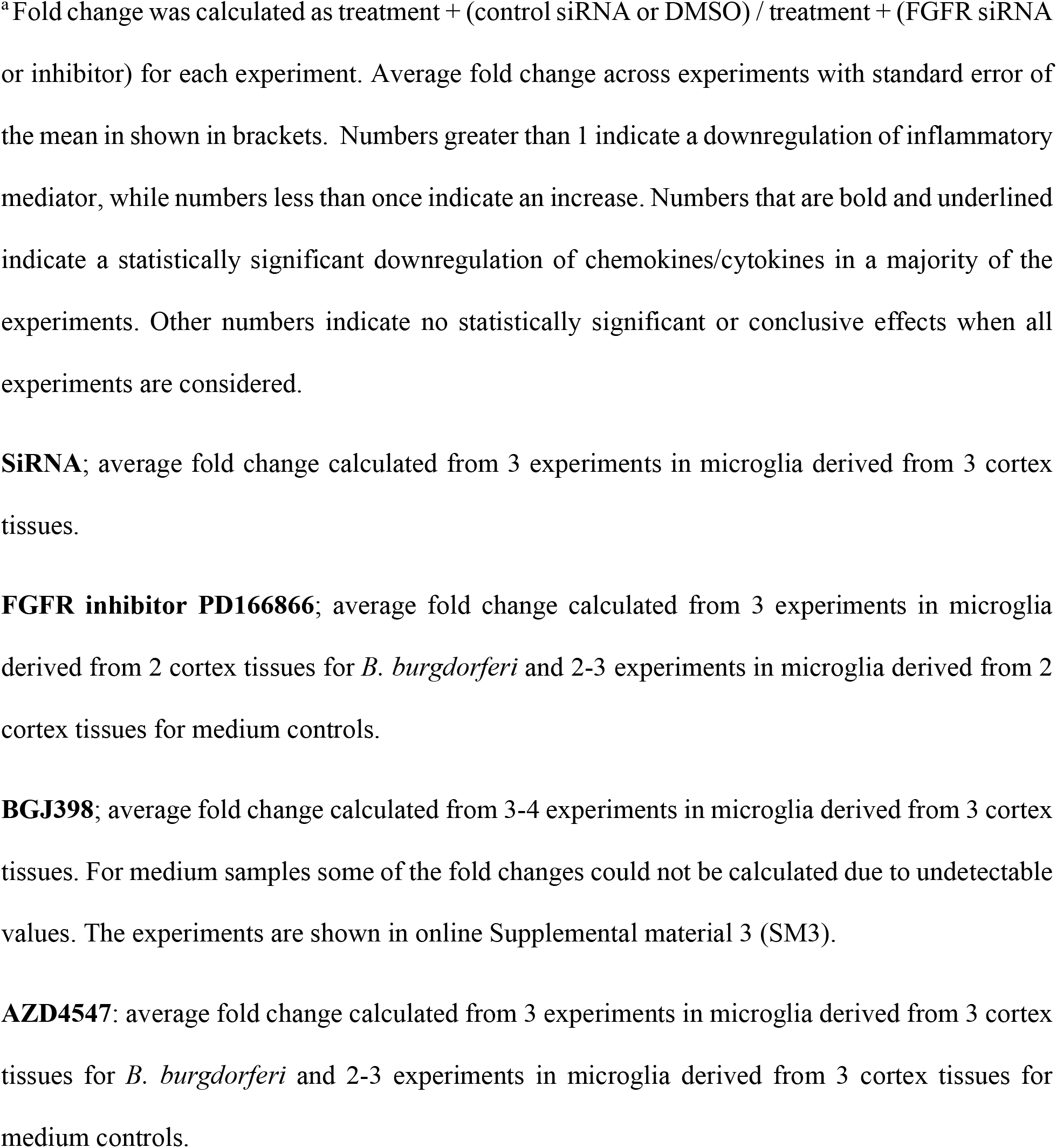
Mean fold downregulation^a^ in inflammatory mediators in response to FGFR inhibition.

| Inhibitor/Treatment |  | IL-6 | IL-8 | MCP-1 |
| --- | --- | --- | --- | --- |
| **Bb 10:1/siRNA** | FGFR1 siRNA (50 nM) | <u>**1.725**</u><br>(±0.565) | <u>**1.151**</u><br>(±0.082) | <u>**1.212**</u><br>(±0.111) |
|  | FGFR2 siRNA (50 nM) | <u>**2.224**</u><br>(±0.916) | <u>**1.214**</u><br>(±0.068) | <u>**1.324**</u><br>(±0.151) |
|  | FGFR3 siRNA (50 nM) | <u>**1.535**</u><br>(±0.325) | <u>**1.274**</u><br>(±0.125) | <u>**1.437**</u><br>(±0.321) |
| **Medium/siRNA** | FGFR1 siRNA (50 nM) | 0.973<br>(±0.112) | 1.073<br>(±0.044) | <u>**1.439**</u><br>(±0.299) |
|  | FGFR2 siRNA (50 nM) | 1.188<br>(±0.070) | 1.194<br>(±0.189) | <u>**1.534**</u><br>(±0.258) |
|  | FGFR3 siRNA (50 nM) | 1.278<br>(±0.165) | <u>**1.571**</u><br>(±0.381) | <u>**1.345**</u><br>(±0.123) |
| **Bb 10:1/FGFR inhibitor PD166866** | 5 µM | <u>**1.982**</u><br>(±0.394) | <u>**1.954**</u><br>(±0.492) | <u>**3.292**</u><br>(±1.056) |
|  | 2.5 µM | <u>**1.850**</u><br>(±0.737) | <u>**1.906**</u><br>(±0.683) | <u>**2.769**</u><br>(±1.397) |
|  | 1 µM | <u>**1.703**</u><br>(±0.474) | <u>**1.713**</u><br>(±0.543) | <u>**2.077**</u><br>(±0.759) |
|  | 500 nM | 1.071<br>(±0.367) | 0.901<br>(±0.226) | 1.113<br>(±0.352) |
| **Medium/FGFR inhibitor PD166866** | 5 µM | 0.880<br>(±0.062) | 1.099<br>(±0.110) | <u>**2.322**</u><br>(±0.615) |
|  | 2.5 µM | 1.133<br>(±0.133) | 1.213<br>(±0.172) | <u>**2.201**</u><br>(±0.417) |
|  | 1 µM | 1.303<br>(±0.048) | <u>**1.217**</u><br>(±0.114) | <u>**1.669**</u><br>(±0.125) |
|  | 500 nM | 1.138<br>(±0.129) | 1.012<br>(±0.092) | <u>**1.402**</u><br>(±0.119) |
| **Bb 10:1/BGJ398** | 1 µM | <u>**2.292**</u><br>(±0.140) | <u>**2.046**</u><br>(±0.158) | <u>**3.480**</u><br>(±0.568) |
|  | 500 nM | <u>**2.403**</u><br>(±0.515) | <u>**1.409**</u><br>(±0.233) | <u>**1.836**</u><br>(±0.301) |
| **Bb 10:1/AZD4547** | 10 µM | <u>**3.817**</u><br>(±1.882) | <u>**2.369**</u><br>(±1.173) | <u>**6.034**</u><br>(±3.967) |
|  | 5 µM | <u>**2.176**</u><br>(±0.921) | <u>**2.048**</u><br>(±0.942) | <u>**2.809**</u><br>(±1.495) |
| **Medium/AZD4547** | 10 µM | 1.424<br>(±0.937) | <u>**1.929**</u><br>(±0.101) | <u>**5.656**</u><br>(±1.626) |
|  | 5 µM | 2.170<br>(±0.581) | <u>**1.338**</u><br>(±0.214) | <u>**2.432**</u><br>(±0.434) |

<sup>a</sup>Fold change was calculated as treatment + (control siRNA or DMSO) / treatment + (FGFR siRNA or inhibitor) for each experiment. Average fold change across experiments with standard error of the mean is shown in brackets. Numbers greater than 1 indicate a downregulation of inflammatory mediator, while numbers less than one indicate an increase. Numbers that are bold and underlined indicate a statistically significant downregulation of chemokines/cytokines in a majority of the experiments. Other numbers indicate no statistically significant or conclusive effects when all experiments are considered.
**SiRNA**; average fold change calculated from 3 experiments in microglia derived from 3 cortex tissues.
**FGFR inhibitor PD166866**; average fold change calculated from 3 experiments in microglia derived from 2 cortex tissues for *B. burgdorferi* and 2-3 experiments in microglia derived from 2 cortex tissues for medium controls.
**BGJ398**; average fold change calculated from 3-4 experiments in microglia derived from 3 cortex tissues. For medium samples some of the fold changes could not be calculated due to undetectable values. The experiments are shown in online Supplemental material 3 (SM3).
**AZD4547**; average fold change calculated from 3 experiments in microglia derived from 3 cortex tissues for *B. burgdorferi* and 2-3 experiments in microglia derived from 3 cortex tissues for medium controls.

In our recent study [28], we showed that non-viable sonicated *B. burgdorferi* can induce inflammation and apoptosis in primary rhesus frontal cortex and dorsal root ganglion tissues. So we next looked at the effect of FGFR1 inhibitor PD166866 on inflammatory mediator output in the presence of sonicated *B. burgdorferi*. The results show that just like its effect on live bacterium-mediated neuroinflammation, the inhibitor also significantly suppressed inflammatory mediators in response to its sonicated contents, implicating novel treatment targets for supplemental therapeutics [Online supplementary material SM3B]. Due to the paucity of primary rhesus microglia availability, and since the FGFR1 receptor alone showed significant efficacy in mediating neuroinflammation, we confined subsequent experiments to this receptor.

### Secreted factors affect FGFR1 expression and signaling

Our next step was to determine what promotes FGFR activation in primary microglia in response to the spirochete. *B. burgdorferi* physically exhibits several TLR ligands but no known ligands that bind FGFRs. Therefore, initial experiments concentrated on whether TLR ligands activate FGFR1 expression. However, preliminary experiments using Pam3CSK4 (Pam3CysSerLys4) and OspA (TLR2), FliC (TLR5) or LPS (TLR4) individually, did not elicit robust expression of FGFR1 as seen with *B. burgdorferi*. Only punctate sporadic expression was generally seen (not shown). It is possible that all three must be simultaneously activated to induce FGFR expression, or dose response studies need to be conducted. Such experiments constitute a study of their own and await tissue availability.

We next looked at whether *B. burgdorferi*-conditioned medium can induce expression of FGFR1 or pFGFR1. As seen in Fig. 4a, supernatants obtained from *B. burgdorferi* exposed cells were able to activate FGFR1 and pFGFR1, while the supernatants obtained from medium alone controls did not, indicating that factors in the supernatants can activate FGFR1. Since FGFs are the likely ligands for FGFR1 activation, we used a custom FGF antibody array as a screen to determine which FGFs are specifically induced. The results are seen in Fig. 4b and online supplemental material SM4. The heat map in Fig. 4b shows upregulation of microglial FGF2, FGF6, FGF10, FGF12 and FGF23 in response to *B. burgdorferi*. In comparison to the other 4 induced FGFs however, whose values were in thousands of Units (Online supplementary material 4, SM4), FGF2 values were in single digits (not shown), and was not considered to be a real upregulation, but an artifact of fold-change. FGF17, 18 and 19 did not show any distinct pattern, while FGFs 4,5,7,8,9,11,13-1B, 16,20,21 and FGF-BP (FGF-binding proteins) showed a distinct downregulation in comparison to medium controls. Though not included in the heatmap, IL-8 and MCP-1 were included as positive controls for the array and their expressions were as expected, validating the array results (SM4). Interestingly, most of the mediators, (with the exception of FGF8, 16,21 and 23) were suppressed by *B. burgdorferi*-induced FGFR1 activation (SM4, and not shown), as their levels went up in the presence of the inhibitor. A principal component analysis of the data showed that *B. burgdorferi*-only group and the medium/Bb +FGFR1 groups segregated as two clusters with an explained variance of 61.9% indicating two separate patterns for the groups (online supplementary material 5 (SM5)). Within each cluster, the *B. burgdorferi*-only group was more spread, indicating the diversity of response to infection from the genetically diverse animals.

**Fig. 4.**
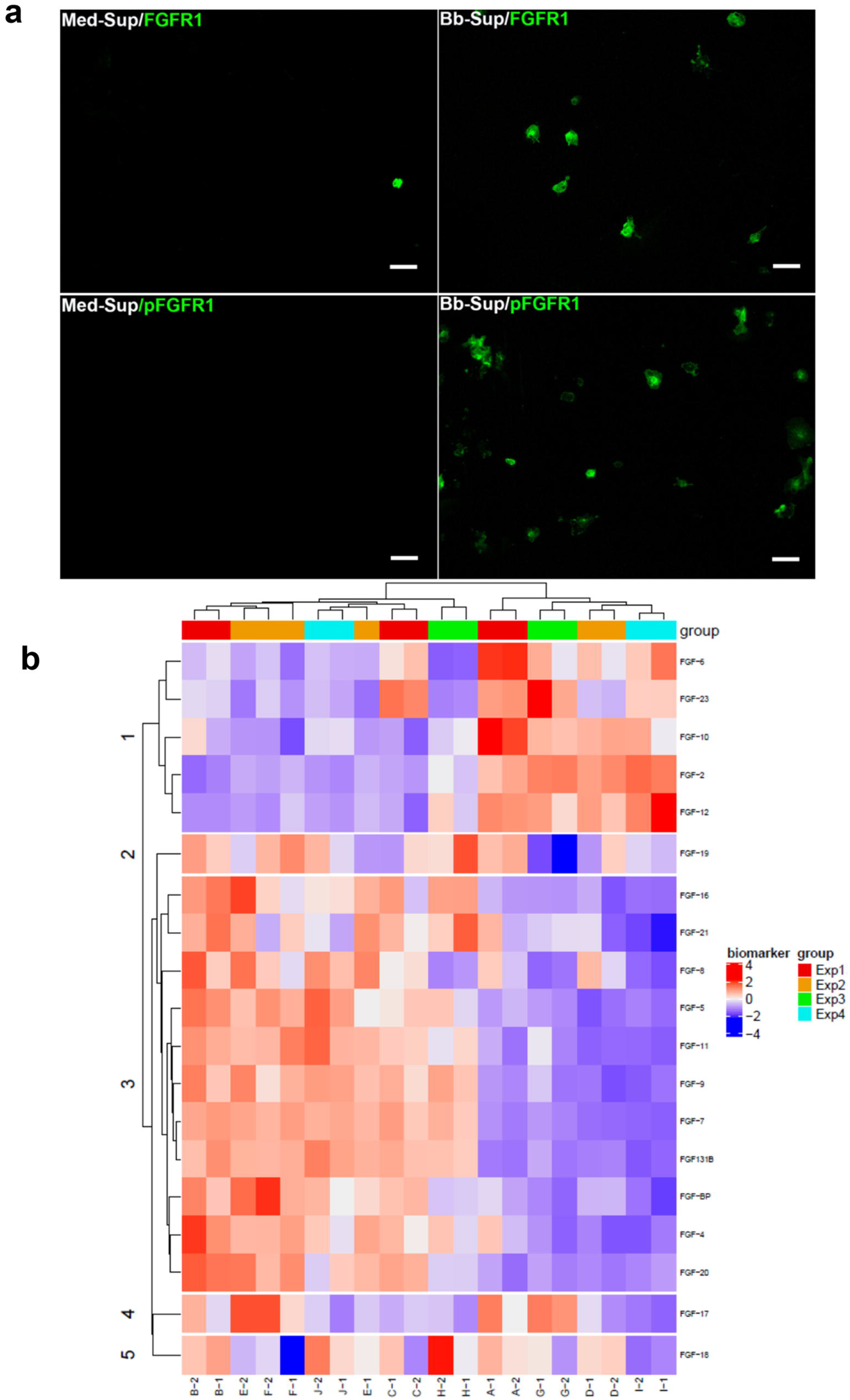
Activation of FGFR1 pathway by microglia conditioned medium (a) and likely FGFs present in the conditioned medium in response to *B. burgdorferi* exposure (b) To test the hypothesis that secreted factors activate the FGFR1 pathway, 24h supernatants from microglia exposed to either *B. burgdorferi* or medium alone were collected, filtered and added to fresh microglial cells from the same tissue for an additional 24 h. Cells were fixed and analyzed for FGFR1 or pFGFR1 by immunofluorescence as before. **a**) shows the activation of FGFR1 or pFGFR1 (both green) by *B. burgdorferi*-exposed microglial conditioned media, indicating activation of this pathway through secreted factors. Bar represents 50 μm. As FGFs are the likely ligands for FGFR1 activation, supernatants from microglia derived from 4 different brain tissues were analyzed for FGF secretion by a custom antibody Array (**b**). Experiment (Exp.) 1: <u>A</u>- Bb 10:1 + DMSO, <u>B</u>-Med + DMSO, <u>C</u>-Bb10:1 + 1 μM PD166866. Exp. 2: <u>D</u>- Bb 10:1 + DMSO, *E*-Med + DMSO, <u>F</u>-Bb10:1 + 1 μM PD166866. Exp. 3: <u>G</u>- Bb 10:1, <u>H</u>- Med. Exp.4: <u>I</u>-Bb 10:1, <u>J</u>- Med.

### Specific FGFs are expressed in primary rhesus microglia in response to *B. burgdorferi*

We next sought to verify some of the antibody array data with additional lines of evidence. As we were mostly interested in factors that likely induced FGFR expression, we focused on the upregulated FGFs. These were FGF6, 10, 12, and FGF23. Fig. 5a shows upregulated expression of FGF6, FGF10 FGF12 and FGF23 in microglial cells in response to *B. burgdorferi* exposure, as verified through immunofluorescence, using antibodies from a different company. Fig.5b shows confocal microscopy of FGF staining in microglial cells, stained for Iba1. Staining was seen along the surface indicating possible engagement with receptors, or at least localized there. Since ELISA assays require a substantial volume of sample materials, we verified only specific FGFs using supernatants from *B. burgdorferi* or medium exposed microglia. FGF6 secretion was additionally verified by ELISA (supplemental material 6 (SM6). FGF12 was only seen by immunofluorescence and not by ELISA. But overall, there was consensus in terms of specificity of induced FGFs with the antibody array. On a technical note, FGF6 was only detected by ELISA when fresh media with fresh serum was used in experiments and detected quickly. Dilution agents also affected detection by ELISA, with PBS being better than standard diluents. The latter reduced detection by approximately 50%. Surprisingly, addition of protease inhibitor phenylmethylsulphonyl fluoride (PMSF) (1mM) lowered the detection levels as well.

**Fig. 5.**
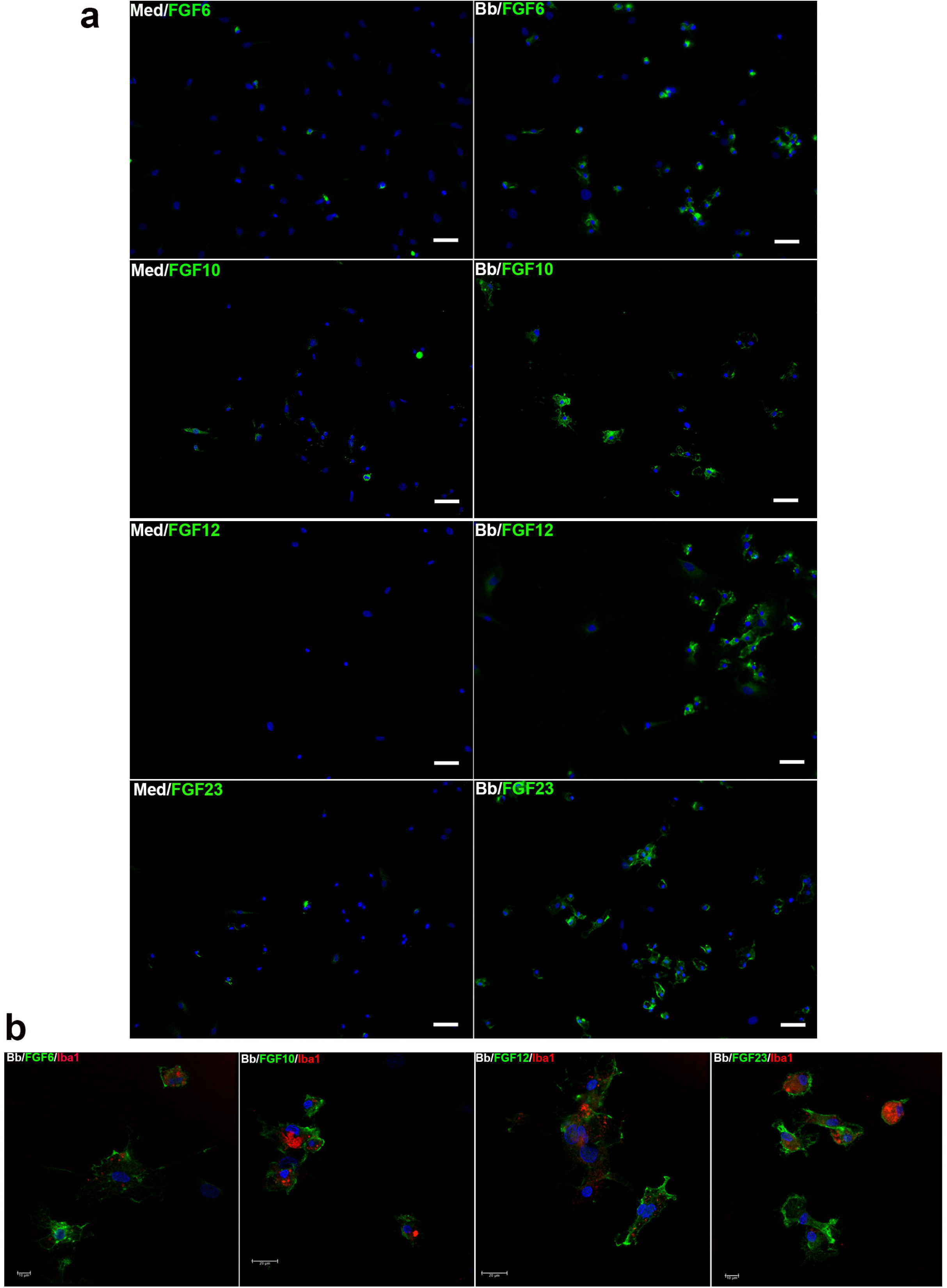
Expression of specific upregulated FGFs in primary rhesus microglia in response to *B. burgdorferi*. The expression of FGF6, FGF10, FGF12 and FGF23 was analyzed by immunofluorescence in primary rhesus microglia. Panel **a** shows immunofluorescent microscopy pictures of the specific FGFs upregulated (green) in response to the Lyme disease bacterium. The nuclei stained with DAPI is shown in blue. Representative pictures from 2 (FGF10, FGF23) to 3 (FGF6, FGF12) experiments are shown. Bar represents 50 μm. Panel **b** shows confocal microscopy pictures of the same FGFs (green) to be microglia specific by staining for Iba1 in red. Nuclear stain is in blue.

### Upregulated FGFs are proinflammatory in a dose dependent manner

We next looked at the role of the induced FGFs in microglial neuroinflammation. We only focused on those FGFs that were also induced through FGFR1, which were FGF6, 10, and 12 (supplemental material 4 (SM4). FGFs were added at various doses on cultured microglial cells for 24 h and supernatants analyzed for IL-6, IL-8 and MCP-1 as before. Results are shown in Fig.6a. FGFs significantly induced production of IL-6 and/or IL-8 but had inconclusive effects on MCP-1 levels (not shown). The effect was also dose dependent and varied with the different FGFs. Increasing doses of FGF6 either increased inflammatory mediator output or remained almost the same. Lower doses of FGF10 (10 ng/ml or 30 ng/ml) did not have any specific effect on cytokine/chemokine levels, but at higher doses (≥50 ng/ml) significantly upregulated IL-8 but not IL-6. FGF12 at lower doses (5 ng/ml) significantly induced IL-6 and IL-8, and also at higher doses (50 ng/ml) but the levels induced were much lower than those induced at 5 ng/ml. This showed that FGFs have proinflammatory effects but in a dose dependent manner. This proinflammatory effect was also seen when FGFs were added in combination. FGF6, 10, and 12 at 5 ng/ml, 20 ng/ml and 5 ng/ml respectively or FGF6, 10, and 12 at 25 ng/ml,60 ng/ml,25 ng/ml respectively, elicited significantly elevated IL-6 and IL-8 compared to controls with no FGFs (not shown).

**Fig. 6.**
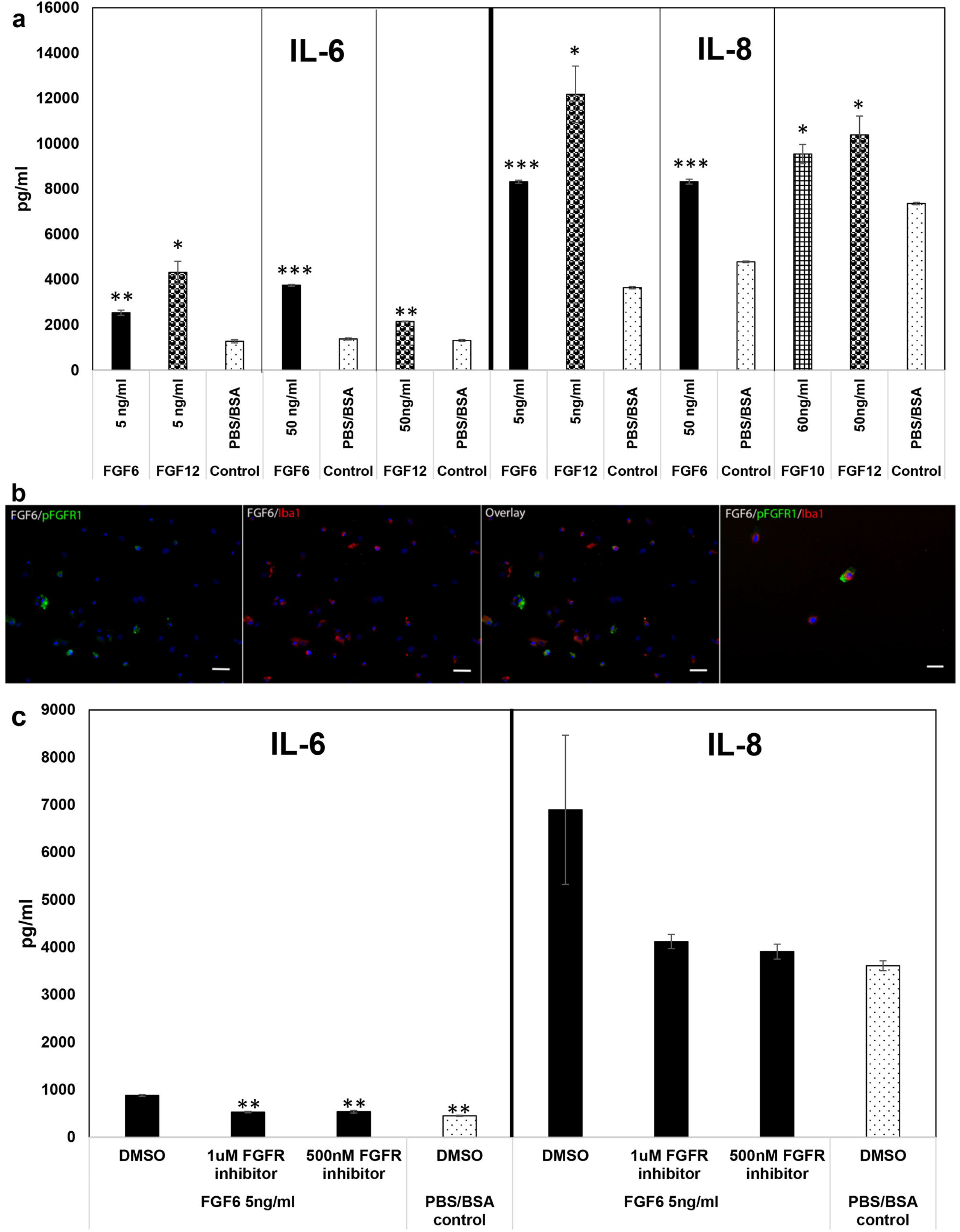
Effect of exogenous addition of FGFs on inflammatory mediator output and activation of FGFR1 pathway on primary rhesus microglia. **a**) Various concentrations of recombinant human FGFs were added to enriched primary rhesus microglial (~80%) cells for 24 h. PBS/BSA (0.1%) was used as the solvent control. Supernatants were collected and analyzed for IL-6, IL-8 and MCP-1 expression by Multiplex ELISA. Lines within each cytokine/chemokine indicates that they were analyzed separately. 5 ng/ml data is representative of 2 experiments conducted on microglia derived from one frontal cortex tissue, while the higher concentration is representative of 2 experiments conducted on microglia derived from 2 different frontal cortex tissues. Data shown are from experiments that were performed with the same animal tissue. Bar represents standard deviation. * p < 0.05, **p < 0.01, and *** p < 0.001. **b**) shows activation of FGFR1 pathway by addition of 5 ng/ml of FGF6 (+ DMSO) to primary rhesus microglial cells. Upregulation of pFGFR1 (green) is seen in cells that also stain for Iba1 (red). Bar represents 50 μm. Panel on the far-right shows the same data at a higher magnification (Bar represents 25 μm). **c**) Shows the effect of PD166866 FGFR1 inhibitor on the inflammatory output in response to exogenous addition of FGF6. A representative experiment is shown of 2-3 experiments carried out on microglia derived from 2 different tissues.

We next confirmed that the effect of FGFs, particularly for FGF6 was through FGFR1. Fig.6b shows upregulation of pFGFR1 in Iba1 stained microglia in response to FGF6, and that inhibition of FGFR1 by PD166866 downregulated the FGF6 mediated upregulation of IL-6 and IL8 (Fig. 6c).

A summary of the data from this study and the proposed model of FGFR activation in primary rhesus microglia is shown in Fig. 7a. Fig. 7b shows the intersectionality of the various FGFs induced or downregulated in microglia in response to live *B. burgdorferi* with other neurological conditions.

**Fig. 7.**
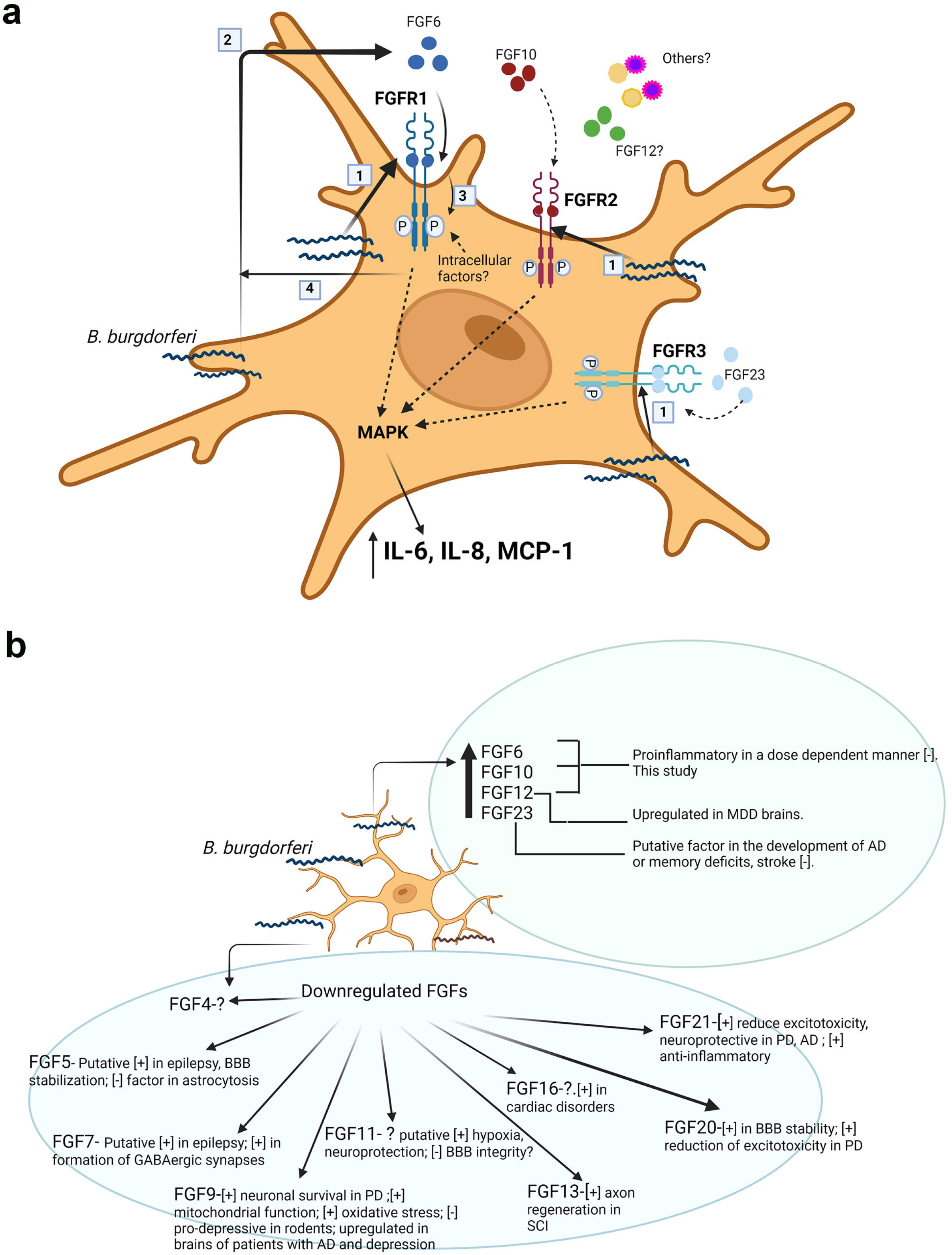
Proposed model of FGFR activation pathways in response to live *B. burgdorferi* exposure (a) and intersectionality of the induced and downregulated FGFs with other neurological conditions (b) **a**). Exposure of primary rhesus microglia to live *B. burgdorferi* upregulates the surface expression of FGFR1, FGFR2 and FGFR3 [1]. Host-pathogen interaction also induces expression of several FGFs such as FGF6, FGF10, FGF12, and FGF23, of which FGF6 (and likely FGF10 and FGF23) are secreted from the cells [2]. Whether FGF12 is secreted extracellularly, is unclear. Ligand binding of FGF6 to FGFR1 induces phosphorylation of the receptor [3] and secretion of IL-6 and IL-8. The intracellular signaling pathway is likely through MAPK pathways, particularly ERK, as have been demonstrated in our previous study in primary rhesus microglia [22]. While FGF6 was shown to activate FGFR1 in this study, it can also activate other FGFRs. Similarly, FGF10, shown to activate FGFR2 in the model, can also activate FGFR1, while FGF23 can activate FGFR3, FGFR2 and FGFR1 [85]. As FGF6, 10 and 12 only activated IL-6 and/or IL-8, but the inhibition of FGFR1 individually by siRNA downregulated IL-6, IL-8 as well as MCP-1, it is likely that other than FGF23, non-FGF molecules present in the supernatant also likely activate this receptor. It should be noted that only autocrine effects of FGF binding FGFRs in microglia are shown. It is possible that some paracrine effects on other glial cells also occur and will be tested in future studies. Finally, our study also demonstrated that synthesis (or inhibition) of FGFs (with the exception of FGF8, 23, and 16 & 21 to an extent) was also through FGFR1(SM4), as suppression of FGFR1 signaling with PD166866 modulated FGF levels [4]. **b**) shows the known neurological roles of the FGFs from this study and others. [-] indicates (putative) negative roles, while [+] indicates (putative) positive effects of the indicated FGFs. The listed roles are not exhaustive. Please see Discussion section for details. Upregulation of FGFs with deleterious effects, and downregulation of FGFs with ameliorative effects can contribute towards Lyme neuroborreliosis sequelae and other neuropathologies. Figures created with BioRender.com

## DISCUSSION

We show in this study using primary rhesus microglia, a rarely available resource, that members of the FGF/FGFR system are novel mediators of pathogenesis in Lyme neuroborreliosis. We show here that FGFRs 1,2 and 3 are activated in response to *B. burgdorferi* (Fig. 1) and that inhibition of these receptors down regulates neuroinflammation (Figs.2, 3, and Table 1). This was determined using multiple lines of evidence including siRNA and three different inhibitors indicating the strength of the data. In our previous study with the oligodendrocyte cell line MO3.13, inhibition with the FGFR1 inhibitor alone (same as used in this study) significantly *<u>increased</u>* chemokines and cytokine levels in direct contrast to the data in primary rhesus microglia [23]. Increase in inflammatory output leads to increased apoptosis in oligodendrocytes [29]. We had postulated in that study, that since MO3.13 oligodendrocyte cell line was created by fusion of oligodendrocytes with rhabdomyosarcoma cells, which overexpress FGFR1 [30], it is likely that the results could be due to this fact. As FGFR inhibitors are being used to treat sarcomas [31], which have various FGFR mutations [32], an increased apoptosis (due to the increased inflammation) would be desired for treatment. Therefore, this result was not surprising in light of that fact, but indicated that this *might be* rhabdomyosarcoma specific effect and not oligodendrocyte mediated. But since the effect was profound, we nevertheless decided to conduct more comprehensive follow-up studies in primary microglial cells without any other confounding factors. And we show here that FGFRs alone do *<u>mediate</u>* neuroinflammation in primary rhesus microglia, and are possible novel mediators of inflammatory pathogenesis in Lyme neuroborreliosis.

We also show that secreted factors in the supernatants activate FGFRs, particularly FGFR1 (Fig. 4). And that several FGFs especially FGF6, secreted into the supernatant can activate the FGFR1 pathway (Fig. 6b and 6c). However, the effect with *B. burgdorferi* supernatant alone was much more profound than that with FGF6 alone (Fig. 5 and Fig.6b) or in combination with other FGFs (FGF10, 12, not shown). It is possible that FGF23, which we did not test, might contribute in activating FGFR1. But it is also likely that non-FGFs activate this pathway. One type of such molecules could be the galectins. Galectins are soluble proteins that contain carbohydrate recognition domains, and play roles in inflammation, signaling and others [33]. While they reside predominantly in the intracellular compartment, they can be secreted by non-classical pathways [34]. In a recent study, extracellular galectins (galectin 1 and 3) have been shown to activate FGFR1 [35], similar to FGFs. Interestingly, in the CNS, galectin 3 is secreted by microglia and is proinflammatory [36]. Other molecules such as bradykinin has been shown to binds its receptor to activate intracellular c-src, which then transactivates FGFR1, independent of FGF-mediated activation [37]. Bradykinin, a peptide, affects blood-brain barrier permeability [38] and is considered a mediator of inflammation, although its neuroprotective roles in rat microglia have also been described [39]. It is not clear if bradykinin is produced in microglia under pathological conditions, but its receptors in microglia have been documented [40]. C-src, a tyrosine kinase, has been demonstrated to activate microglia and is considered proinflammatory in many studies [41, 42]. It is also possible that even without bradykinin release, c-src alone could transactivate FGFR1 receptor, without ligand binding.

One other interesting observation we made in this study was that inhibition of FGFRs in medium alone controls downregulated MCP-1 production with all the inhibitors, be it siRNA or others. This indicated that some low level activation and signaling (pFGFR1) is on, that is not greatly detected by immunofluorescence. But a hint of this effect could be seen in Fig.1a, where some receptor expression is seen. The only likely component in the medium that could elicit this activation is the FBS. Incidentally, serum is associated with galectin 3 secretion as its secretion was shown to be decreased in serum-free media [43]. In summary, it is possible that upon *B. burgdorferi* addition, cumulative activation of several TLRs by *B. burgdorferi* ligands causes increased levels of all these molecules (galectins, bradykinin or c-src) along with increased protein expression of FGFRs and surface expression. Increased activation of the FGFR receptors and subsequent signaling causes increased production of specific FGFs while downregulating other FGFs. This also sets up an autocrine loop to ensue, where the induced FGFs likely cause their own production subsequently. Low level activation of FGFR1 in medium alone would induce low levels of MCP-1, while increased activation causes upregulation of specific FGFs such as FGF6 and others, which in-turn contribute to IL6 and IL-8 levels. An overview of the data obtained from this study is shown in Fig.7a.

With respect to roles of individual FGFs, FGF6, FGF10, FGF12, and FGF23 were shown to be upregulated in microglia in response to *B. burgdorferi*. FGF6 has been shown to be associated with muscle growth [44], but not many studies exist with regard to CNS. One study showed that it is involved in brain development in the late embryonic stages [45], while another showed that human umbilical mesenchymal stem cells secrete FGF6 among others, and that transplantation of stem cells in epileptic rats downregulated microglial activation. Whether this effect was mediated by FGF6 or others is not clear [46]. In another study, FGF6 was demonstrated to be secreted in human fetal astrocytes, and treatment with alpha-synuclein decreased its levels after 48h [47]. However, no role for this cytokine has been demonstrated. We show here for the first time that infection can trigger FGF6 in microglia and it is proinflammatory [Figs. 4b, 5, 6 and SM6]. <u>To our knowledge this is the first report to define a role for FGF6.</u>

FGF10 has generally been shown to be neuroprotective in rodent models of spinal cord injury, neuroinflammation and others, both *in vitro* in BV2 microglial cells and *in vivo* in mice/rats [48–50]. In our study using rhesus microglia, we show that the FGF10 effect is dose dependent. Lower doses did not have any significant effect, while higher doses (≥ 50 ng/ml) significantly induced IL-8, but had no significant effect on IL-6. Since 100 ng/ml was used in BV2 cells, it is not clear if its protective at very high doses, or if its due to species difference, or cell line effects. It should be noted that deletion of an FGF gene may result in a completely different phenotype than when its levels are modulated. Therefore, functions attributed to FGFs due to gene deletions may or may not reflect disease pathogenesis where its levels can vary.

FGF12 gene expression has been shown to be induced in BV2 microglia in response to LPS [51], however <u>no role for it has been described until this study.</u> The main focus of FGF12 studies has been with its genetic alterations and associated epileptic changes through its ability to bind voltage-gated sodium ion channels [52, 53]. Its expression has also been shown to be elevated in anterior cingulate cortex of patients with major depressive disorder (MDD) [54]. However, an interesting anomaly is that FGF12 (along with FGF11,13, and 14) is considered intracellular. A pioneering study in human embryonic kidney cells (HEK 293, epithelial morphology) showed that transfection of the cells with *FGF12* gene caused accumulation of the protein in the nucleus with no detectable secretion [55]. In our study, microglial supernatants were analyzed by the antibody array and FGF12 was found to be elevated in the extracellular environment. Confocal microscopy also showed likely surface location of FGF12 in microglia (Fig. 5b). However, we could not detect FGF12 in the supernatants by ELISA. So we cannot confirm whether FGF12 is actually secreted outside, like interleukins. It is possible it is secreted but surface located and not truly in the extracellular environment. Our hypotheses for this anomaly between array and ELISA are 1) As the array is much more sensitive than ELISA, intracellular FGF12 was detected by the array due to possible breach of cellular contents, or presence of some cells in the supernatants. 2) FGF12 is secreted, and as the array procedure biotinylates the proteins prior to detection, FGFs are stabilized and better detected. In ELISA assays due to long incubations prior to detection, the natural confirmation destabilizes quickly and hence is not detected. The latter hypothesis could be tested by ELISA of the cell lysates, but await tissue availability. More studies in other primary cell types are needed to clarify this issue, and not just immortalized cells. The intracellular class of FGFs or FGF homologous factors (FHFs) as they are known, were also thought to not activate FGFRs [56]. Recent studies shown that is not the case [57] and in our study cells did respond to exogenous addition of FGF12 in inducing IL-6 and IL-8, and is *possibly* through FGFRs as well. Therefore, the characterization of these factors is far from complete. <u>This is also the first study (to our knowledge) to show a role for FGF12 in the brain glial cells.</u>

FGF23, an endocrine hormone secreted by osteocytes is required for maintaining phosphate homeostasis. Due to this function, it has long known for its role in chronic kidney disease, characterized by elevated FGF23 levels and hyperphosphatemia. Outside the kidney, its role in CNS have also been delineated. Mice overexpressing FGF23 had impaired spatial memory and learning [58], and other studies show that exogenous FGF23 can reduce proximal arborization in hippocampal neurons, impacting memory functions [59]. Recent studies in patients show that high levels of FGF23 in the serum is associated with risk for stroke [60] and dementia [61]. <u>Thus, the FGFs upregulated by *B. burgdorferi* in microglia are likely deleterious in the long run.</u>

With respect to the other FGFs that were downregulated in response to *B. burgdorferi* exposure, only some of the salient ones will be discussed here. In terms of modulating neuroinflammatory mediators *per se*, not much data exist for much of the FGFs. FGF20 was shown to be protective in blood brain barrier disruption by upregulating tight junction proteins, increasing the transelectrical endothelial resistance and reducing neuroinflammation in traumatic brain injury models [62]. FGF21 is the most studied in term of neuroinflammation and almost all describe an anti-inflammatory role. FGF21 administration was shown to protect against neuroinflammation in oxidative stress, ischemic stroke, and in obesity [63–65]. In terms of other neurological conditions, FGF4 expression was upregulated in patients’ CSF transitioning from mild cognitive impairment to AD progression [66]. Its role, however is not known. FGF5 expression was shown to be elevated in astrocytic tumors implying a role in astrogliosis [67]. Deletion of this gene and *Fgf2* in mice caused increased BBB permeability [68]. As BBB leakage can correlate with epilepsy, it is likely a positive factor in preventing seizures [69]. Similar to FGF5, FGF7 also has a putative positive role in epilepsy as *Fgf7*-deficient mice exhibit enhanced seizure activity [70]. Clustering of GABAergic synaptic vesicles was also reduced in *Fgf7* deleted mice, implying a role in GABAergic synapse formation. Incidentally, low GABA levels can cause depression, anxiety and others [71, 72]. FGF9 immunoreactivity was demonstrated in the brains of AD patients and those with MDD [54, 73]. It has protective roles in PD by downregulating oxidative stress, improving mitochondrial function and promoting neuronal survival [14]. Contrarily, it had pro-anxiety and depressive effects as exogenous administration increased these behaviors in rats [74]. Hypoxia inducible factor-1α (HIF-1α) is a transcription factor required for cellular adaption to hypoxia. FGF11 level was shown to be increased in hypoxic conditions and was demonstrated to stabilize HIF-1α [75]. However, HIF-1α has contrary roles in neuroprotection [76], and blood brain barrier disruption [77], so the role of FGF11 is unclear. In a rat model of spinal cord injury, FGF13 was demonstrated to promote axon regeneration, by stabilizing microtubules and promoting mitochondria function [78]. FGF16 was shown to provide cardiac protection in diabetes after myocardial infarction [79]. Other than brain development [80] its role in neurological conditions is unknown. FGF20 protected dopaminergic neurons in the substantia nigra in a rat model of PD [81], likely by reducing excitotoxicity and promoting survival [14]. FGF21 has similarly been demonstrated to reduce excitotoxicity, reduce α-synuclein and promote survival of dopaminergic neurons in PD models [14]. Similar protective effects of FGF21 in several *in vitro* and *in vivo* AD models have also been described [82–84]. A summary of these effects is depicted in Fig.7b. <u>By suppressing ameliorative FGFs, *B. burgdorferi* infection likely accelerates underlying comorbidities and hastens manifestations.</u>

### Concluding remarks

*B. burgdorferi* infection has been shown to induce psychiatric changes, secondary dementia, anxiety and depression in human patients. The ability of the bacterium to induce pathogenic FGFs involved in depression and memory deficits and downregulate protective FGFs that can alleviate several neurological conditions suggests that the FGF system likely lies at the intersection of Lyme neuroborreliosis sequelae and other neurological conditions. Presence of Lyme infection in case reports with PD, AD, Lewy Body disease and others suggests that a chronic infection with *B. burgdorferi* can exacerbate or accelerate pathology in susceptible individuals with underlying comorbidities through an FGF/FGFR mediated process. It can also complicate treatment modalities. Whether *B. burgdorferi* alone can cause complex multifactorial diseases such as AD or PD is unclear and remains to be tested using single factorial approach in appropriate animal models. As microglia share similar functionality with macrophages, we expect similar FGF modulations in the periphery also. Since this study utilized a single glial cell type to study FGF/FGFR system, we hope to conduct follow-up studies *in vivo* in relevant animal models. We also hope to assess the FGF system in human Lyme disease patients to correlate specific FGFs with symptomology. As FGFR1 also contributed towards neuroinflammation mediated by non-viable *B. burgdorferi*, it poses an attractive target for anti-inflammatory treatments in antibiotic refractive conditions. In conclusion, in this study we show a novel molecular basis for neuroinflammation associated with Lyme neuroborreliosis and its sequelae.

## LIST OF ABBREVIATIONS

AD: Alzheimer’s disease
ALS: amyotrophic lateral sclerosis
BBB: blood brain barrier
BSA: bovine serum albumin
CNS: central nervous system
DMSO: dimethyl sulfoxide
EAE: experimental autoimmune encephalitis
ELISA: enzyme linked immunosorbent assays
FBS: fetal bovine serum
FGF: fibroblast growth factor
FGFR: fibroblast growth factor receptor
FTD: fronto temporal dementia
GABA: gamma aminobutyric acid
GM-CSF: granulocyte macrophage colony stimulating factor
HIF-1α: Hypoxia inducible factor-1α
IL-6: interleukin 6
IL-8: interleukin 8
LBD: Lewy Body Dementia
LD: Lyme disease
LNB: Lyme neuroborreliosis
LPS: lipopolysaccharide
MCP-1: monocyte chemoattractant protein-1
MDD: major depressive disorder
MOI: multiplicity of infection
NGS: normal goat serum
PBS: phosphate buffered saline
PD: Parkinson’s disease
PMSF: phenylmethylsulphonyl fluoride
PNS: peripheral nervous system
RNAi: ribonucleic acid interference
siRNA: small interfering ribonucleic acid
TLR-Toll: like receptor.

## DECLARATIONS

### AVAILABILITY OF DATA AND MATERIALS

All the data are in the manuscript or in the supplemental data. Any other datasets used and/or analyzed during the current study are available from the corresponding author on reasonable request

### CONSENT FOR PUBLICATION

Not Applicable

### ETHICS APPROVAL

No experiments were conducted on live animals. Euthanasia procedures were carried out according to the Tulane Institutional Animal Care and Use Committee (Tulane IACUC) guidelines.

### AUTHOR CONTRIBUTIONS

GP conceived the study, performed all the experiments, individual ELISAs, immunofluorescence, analyzed the data and wrote the manuscript. MBP performed the multiplex ELISAs and CCM did the confocal microscopy.

## ACKNOWLEDGEMENTS

We thank Mr. Maury Duplantis and Dr. Robert Blair, TNPRC, for help with tissue availability. We thank Ms. Mary Barnes and Dr. Carolina Allers Hernandez, TNPRC, for help with the Multiplex assays. We thank Dr. Kelly Whittaker and technical staff at RayBiotech services, for help with the antibody array. We thank Dr. Mario Philipp (Ret.), former Chair, Bacteriology and Parasitology, TNPRC, for manuscript edits.

## REFERENCES

1. Rodino KG, Theel ES, Pritt BS: Tick-Borne Diseases in the United States. Clin Chem 2020, 66:537–548.

2. Kugeler KJ, Schwartz AM, Delorey MJ, Mead PS, Hinckley AF: Estimating the Frequency of Lyme Disease Diagnoses, United States, 2010-2018. Emerg Infect Dis 2021, 27:616–619.

3. Mead PS: Epidemiology of Lyme disease. Infect Dis Clin North Am 2015, 29:187–210.

4. Fallon BA, Nields JA: Lyme disease: a neuropsychiatric illness. Am J Psychiatry 1994, 151:1571–1583.

5. Kristoferitsch W, Aboulenein-Djamshidian F, Jecel J, Rauschka H, Rainer M, Stanek G, Fischer P: Secondary dementia due to Lyme neuroborreliosis. Wien Klin Wochenschr 2018, 130:468–478.

6. MacDonald AB: Borrelia in the brains of patients dying with dementia. JAMA 1986, 256:2195–2196.

7. MacDonald AB, Miranda JM: Concurrent neocortical borreliosis and Alzheimer’s disease. Hum Pathol 1987, 18:759–761.

8. Cassarino DS, Quezado MM, Ghatak NR, Duray PH: Lyme-associated parkinsonism: a neuropathologic case study and review of the literature. Arch Pathol Lab Med 2003, 127:1204–1206.

9. Waniek C, Prohovnik I, Kaufman MA, Dwork AJ: Rapidly progressive frontal-type dementia associated with Lyme disease. J Neuropsychiatry Clin Neurosci 1995, 7:345–347.

10. Gadila SKG, Rosoklija G, Dwork AJ, Fallon BA, Embers ME: Detecting Borrelia Spirochetes: A Case Study With Validation Among Autopsy Specimens. Front Neurol 2021, 12:628045.

11. Paradiso B, Zucchini S, Simonato M: Implication of fibroblast growth factors in epileptogenesis-associated circuit rearrangements. Front Cell Neurosci 2013, 7:152.

12. Terwisscha van Scheltinga AF, Bakker SC, Kahn RS: Fibroblast growth factors in schizophrenia. Schizophr Bull 2010, 36:1157–1166.

13. Alam R, Mrad Y, Hammoud H, Saker Z, Fares Y, Estephan E, Bahmad HF, Harati H, Nabha S: New insights into the role of fibroblast growth factors in Alzheimer’s disease. Mol Biol Rep 2022, 49:1413–1427.

14. Liu Y, Deng J, Liu Y, Li W, Nie X: FGF, Mechanism of Action, Role in Parkinson’s Disease, and Therapeutics. Front Pharmacol 2021, 12:675725.

15. Deng Z, Deng S, Zhang MR, Tang MM: Fibroblast Growth Factors in Depression. Front Pharmacol 2019, 10:60.

16. Dordoe C, Chen K, Huang W, Chen J, Hu J, Wang X, Lin L: Roles of Fibroblast Growth Factors and Their Therapeutic Potential in Treatment of Ischemic Stroke. Front Pharmacol 2021, 12:671131.

17. Rajendran R, Bottiger G, Stadelmann C, Karnati S, Berghoff M: FGF/FGFR Pathways in Multiple Sclerosis and in Its Disease Models. Cells 2021, 10.

18. Woodbury ME, Ikezu T: Fibroblast growth factor-2 signaling in neurogenesis and neurodegeneration. J Neuroimmune Pharmacol 2014, 9: 92–101.

19. Li S, Bock E, Berezin V: Neuritogenic and neuroprotective properties of peptide agonists of the fibroblast growth factor receptor. Int J Mol Sci 2010, 11:2291–2305.

20. Cassina P, Pehar M, Vargas MR, Castellanos R, Barbeito AG, Estevez AG, Thompson JA, Beckman JS, Barbeito L: Astrocyte activation by fibroblast growth factor-1 and motor neuron apoptosis: implications for amyotrophic lateral sclerosis. J Neurochem 2005, 93:38–46.

21. von Bartheld CS, Bahney J, Herculano-Houzel S: The search for true numbers of neurons and glial cells in the human brain: A review of 150 years of cell counting. J Comp Neurol 2016, 524:3865–3895.

22. Parthasarathy G, Philipp MT: Inflammatory mediator release from primary rhesus microglia in response to Borrelia burgdorferi results from the activation of several receptors and pathways. J Neuroinflammation 2015, 12:60.

23. Parthasarathy G, Philipp MT: Receptor tyrosine kinases play a significant role in human oligodendrocyte inflammation and cell death associated with the Lyme disease bacterium Borrelia burgdorferi. J Neuroinflammation 2017, 14:110.

24. Mohammadi M, Schlessinger J, Hubbard SR: Structure of the FGF receptor tyrosine kinase domain reveals a novel autoinhibitory mechanism. Cell 1996, 86:577–587.

25. Chell V, Balmanno K, Little AS, Wilson M, Andrews S, Blockley L, Hampson M, Gavine PR, Cook SJ: Tumour cell responses to new fibroblast growth factor receptor tyrosine kinase inhibitors and identification of a gatekeeper mutation in FGFR3 as a mechanism of acquired resistance. Oncogene 2013, 32:3059–3070.

26. Komla-Ebri D, Dambroise E, Kramer I, Benoist-Lasselin C, Kaci N, Le Gall C, Martin L, Busca P, Barbault F, Graus-Porta D, et al: Tyrosine kinase inhibitor NVP-BGJ398 functionally improves FGFR3-related dwarfism in mouse model. J Clin Invest 2016, 126:1871–1884.

27. Risuleo G, Ciacciarelli M, Castelli M, Galati G: The synthetic inhibitor of fibroblast growth factor receptor PD166866 controls negatively the growth of tumor cells in culture. J Exp Clin Cancer Res 2009, 28:151.

28. Parthasarathy G, Gadila SKG: Neuropathogenicity of non-viable Borrelia burgdorferi ex vivo. Sci Rep 2022, 12:688.

29. Ramesh G, Benge S, Pahar B, Philipp MT: A possible role for inflammation in mediating apoptosis of oligodendrocytes as induced by the Lyme disease spirochete Borrelia burgdorferi. J Neuroinflammation 2012, 9:72.

30. Goldstein M, Meller I, Orr-Urtreger A: FGFR1 over-expression in primary rhabdomyosarcoma tumors is associated with hypomethylation of a 5’ CpG island and abnormal expression of the AKT1, NOG, and BMP4 genes. Genes Chromosomes Cancer 2007, 46:1028–1038.

31. Chudasama P, Renner M, Straub M, Mughal SS, Hutter B, Kosaloglu Z, Schwessinger R, Scheffler M, Alldinger I, Schimmack S, et al: Targeting Fibroblast Growth Factor Receptor 1 for Treatment of Soft-Tissue Sarcoma. Clin Cancer Res 2017, 23:962–973.

32. Napolitano A, Ostler AE, Jones RL, Huang PH: Fibroblast Growth Factor Receptor (FGFR) Signaling in GIST and Soft Tissue Sarcomas. Cells 2021, 10.

33. Johannes L, Jacob R, Leffler H: Galectins at a glance. J Cell Sci 2018, 131.

34. Popa SJ, Stewart SE, Moreau K: Unconventional secretion of annexins and galectins. Semin Cell Dev Biol 2018, 83:42–50.

35. Kucinska M, Porebska N, Lampart A, Latko M, Knapik A, Zakrzewska M, Otlewski J, Opalinski L: Differential regulation of fibroblast growth factor receptor 1 trafficking and function by extracellular galectins. Cell Commun Signal 2019, 17:65.

36. Siew JJ, Chen HM, Chen HY, Chen HL, Chen CM, Soong BW, Wu YR, Chang CP, Chan YC, Lin CH, et al: Galectin-3 is required for the microglia-mediated brain inflammation in a model of Huntington’s disease. Nat Commun 2019, 10:3473.

37. Terzuoli E, Corti F, Nannelli G, Giachetti A, Donnini S, Ziche M: Bradykinin B2 Receptor Contributes to Inflammatory Responses in Human Endothelial Cells by the Transactivation of the Fibroblast Growth Factor Receptor FGFR-1. Int J Mol Sci 2018, 19.

38. Abbott NJ: Inflammatory mediators and modulation of blood-brain barrier permeability. Cell Mol Neurobiol 2000, 20:131–147.

39. Noda M, Kariura Y, Pannasch U, Nishikawa K, Wang L, Seike T, Ifuku M, Kosai Y, Wang B, Nolte C, et al: Neuroprotective role of bradykinin because of the attenuation of pro-inflammatory cytokine release from activated microglia. J Neurochem 2007, 101:397–410.

40. Noda M, Kariura Y, Amano T, Manago Y, Nishikawa K, Aoki S, Wada K: Expression and function of bradykinin receptors in microglia. Life Sci 2003, 72:1573–1581.

41. Socodato R, Portugal CC, Domith I, Oliveira NA, Coreixas VS, Loiola EC, Martins T, Santiago AR, Paes-de-Carvalho R, Ambrosio AF, Relvas JB: c-Src function is necessary and sufficient for triggering microglial cell activation. Glia 2015, 63:497–511.

42. Yang H, Wang L, Zang C, Wang Y, Shang J, Zhang Z, Liu H, Bao X, Wang X, Zhang D: Src Inhibition Attenuates Neuroinflammation and Protects Dopaminergic Neurons in Parkinson’s Disease Models. Front Neurosci 2020, 14:45.

43. Sato S, Burdett I, Hughes RC: Secretion of the baby hamster kidney 30-kDa galactose-binding lectin from polarized and nonpolarized cells: a pathway independent of the endoplasmic reticulum-Golgi complex. Exp Cell Res 1993, 207:8–18.

44. Xu Y, Tan Q, Hu P, Yao J: Characterization and expression analysis of FGF6 (fibroblast growth factor 6) genes of grass carp (Ctenopharyngodon idellus) reveal their regulation on muscle growth. Fish Physiol Biochem 2019, 45:1649–1662.

45. Ozawa K, Uruno T, Miyakawa K, Seo M, Imamura T: Expression of the fibroblast growth factor family and their receptor family genes during mouse brain development. Brain Res Mol Brain Res 1996, 41:279–288.

46. Huang PY, Shih YH, Tseng YJ, Ko TL, Fu YS, Lin YY: Xenograft of human umbilical mesenchymal stem cells from Wharton’s jelly as a potential therapy for rat pilocarpine-induced epilepsy. Brain Behav Immun 2016, 54:45–58.

47. Sengul B, Dursun E, Verkhratsky A, Gezen-Ak D: Overexpression of alpha-Synuclein Reorganises Growth Factor Profile of Human Astrocytes. Mol Neurobiol 2021, 58:184–203.

48. Chen J, Wang Z, Zheng Z, Chen Y, Khor S, Shi K, He Z, Wang Q, Zhao Y, Zhang H, et al: Neuron and microglia/macrophage-derived FGF10 activate neuronal FGFR2/PI3K/Akt signaling and inhibit microglia/macrophages TLR4/NF-kappaB-dependent neuroinflammation to improve functional recovery after spinal cord injury. Cell Death Dis 2017, 8:e3090.

49. Fang M, Jiang S, Zhu J, Fu X, Hu Y, Pan S, Jiang H, Lin J, Yuan J, Li P, Lin Z: Protective effects of FGF10 on neurovascular unit in a rat model of neonatal hypoxic-ischemic brain injury. Exp Neurol 2020, 332:113393.

50. Li YH, Fu HL, Tian ML, Wang YQ, Chen W, Cai LL, Zhou XH, Yuan HB: Neuron-derived FGF10 ameliorates cerebral ischemia injury via inhibiting NF-kappaB-dependent neuroinflammation and activating PI3K/Akt survival signaling pathway in mice. Sci Rep 2016, 6:19869.

51. Sohn SH, Chung HS, Ko E, Jeong HJ, Kim SH, Jeong JH, Kim Y, Shin M, Hong M, Bae H: The genome-wide expression profile of Nelumbinis semen on lipopolysaccharide-stimulated BV-2 microglial cells. Biol Pharm Bull 2009, 32:1012–1020.

52. Veliskova J, Marra C, Liu Y, Shekhar A, Park DS, Iatckova V, Xie Y, Fishman GI, Velisek L, Goldfarb M: Early onset epilepsy and sudden unexpected death in epilepsy with cardiac arrhythmia in mice carrying the early infantile epileptic encephalopathy 47 gain-of-function FHF1(FGF12) missense mutation. Epilepsia 2021, 62:1546–1558.

53. Willemsen MH, Goel H, Verhoeven JS, Braakman HMH, de Leeuw N, Freeth A, Minassian BA: Epilepsy phenotype in individuals with chromosomal duplication encompassing FGF12. Epilepsia Open 2020, 5:301–306.

54. Evans SJ, Choudary PV, Neal CR, Li JZ, Vawter MP, Tomita H, Lopez JF, Thompson RC, Meng F, Stead JD, et al: Dysregulation of the fibroblast growth factor system in major depression. Proc Natl Acad Sci U S A 2004, 101:15506–15511.

55. Smallwood PM, Munoz-Sanjuan I, Tong P, Macke JP, Hendry SH, Gilbert DJ, Copeland NG, Jenkins NA, Nathans J: Fibroblast growth factor (FGF) homologous factors: new members of the FGF family implicated in nervous system development. Proc Natl Acad Sci U S A 1996, 93:9850–9857.

56. Olsen SK, Garbi M, Zampieri N, Eliseenkova AV, Ornitz DM, Goldfarb M, Mohammadi M: Fibroblast growth factor (FGF) homologous factors share structural but not functional homology with FGFs. J Biol Chem 2003, 278:34226–34236.

57. Sochacka M, Opalinski L, Szymczyk J, Zimoch MB, Czyrek A, Krowarsch D, Otlewski J, Zakrzewska M: FHF1 is a bona fide fibroblast growth factor that activates cellular signaling in FGFR-dependent manner. Cell Commun Signal 2020, 18:69.

58. Liu P, Chen L, Bai X, Karaplis A, Miao D, Gu N: Impairment of spatial learning and memory in transgenic mice overexpressing human fibroblast growth factor-23. Brain Res 2011, 1412:9–17.

59. Hensel N, Schon A, Konen T, Lubben V, Forthmann B, Baron O, Grothe C, Leifheit-Nestler M, Claus P, Haffner D: Fibroblast growth factor 23 signaling in hippocampal cells: impact on neuronal morphology and synaptic density. J Neurochem 2016, 137:756–769.

60. Yao XY, Li S, Zhang LG, Liu ZH, Bao JN, Wu ZY: Higher Serum Fibroblast Growth Factor-23 Levels and the Risk of Stroke and Its Subtypes: Evidence From a Meta-Analysis of Prospective Studies. J Stroke Cerebrovasc Dis 2018, 27:3076–3083.

61. McGrath ER, Himali JJ, Levy D, Conner SC, Pase MP, Abraham CR, Courchesne P, Satizabal CL, Vasan RS, Beiser AS, Seshadri S: Circulating fibroblast growth factor 23 levels and incident dementia: The Framingham heart study. PLoS One 2019, 14:e0213321.

62. Chen J, Wang X, Hu J, Du J, Dordoe C, Zhou Q, Huang W, Guo R, Han F, Guo K, et al: FGF20 Protected Against BBB Disruption After Traumatic Brain Injury by Upregulating Junction Protein Expression and Inhibiting the Inflammatory Response. Front Pharmacol 2020, 11:590669.

63. Kang K, Xu P, Wang M, Chunyu J, Sun X, Ren G, Xiao W, Li D: FGF21 attenuates neurodegeneration through modulating neuroinflammation and oxidant-stress. Biomed Pharmacother 2020, 129:110439.

64. Wang D, Liu F, Zhu L, Lin P, Han F, Wang X, Tan X, Lin L, Xiong Y: FGF21 alleviates neuroinflammation following ischemic stroke by modulating the temporal and spatial dynamics of microglia/macrophages. J Neuroinflammation 2020, 17:257.

65. Wang Q, Yuan J, Yu Z, Lin L, Jiang Y, Cao Z, Zhuang P, Whalen MJ, Song B, Wang XJ, et al: FGF21 Attenuates High-Fat Diet-Induced Cognitive Impairment via Metabolic Regulation and Anti-inflammation of Obese Mice. Mol Neurobiol 2018, 55:4702–4717.

66. Lehallier B, Essioux L, Gayan J, Alexandridis R, Nikolcheva T, Wyss-Coray T, Britschgi M, Alzheimer’s Disease Neuroimaging I: Combined Plasma and Cerebrospinal Fluid Signature for the Prediction of Midterm Progression From Mild Cognitive Impairment to Alzheimer Disease. JAMA Neurol 2016, 73:203–212.

67. Allerstorfer S, Sonvilla G, Fischer H, Spiegl-Kreinecker S, Gauglhofer C, Setinek U, Czech T, Marosi C, Buchroithner J, Pichler J, et al: FGF5 as an oncogenic factor in human glioblastoma multiforme: sautocrine and paracrine activities. Oncogene 2008, 27:4180–4190.

68. Reuss B, Dono R, Unsicker K: Functions of fibroblast growth factor (FGF)-2 and FGF-5 in astroglial differentiation and blood-brain barrier permeability: evidence from mouse mutants. J Neurosci 2003, 23:6404–6412.

69. van Vliet EA, da Costa Araujo S, Redeker S, van Schaik R, Aronica E, Gorter JA: Blood-brain barrier leakage may lead to progression of temporal lobe epilepsy. Brain 2007, 130:521–534.

70. Terauchi A, Johnson-Venkatesh EM, Toth AB, Javed D, Sutton MA, Umemori H: Distinct FGFs promote differentiation of excitatory and inhibitory synapses. Nature 2010, 465:783–787.

71. Schur RR, Draisma LW, Wijnen JP, Boks MP, Koevoets MG, Joels M, Klomp DW, Kahn RS, Vinkers CH: Brain GABA levels across psychiatric disorders: A systematic literature review and meta-analysis of (1) H-MRS studies. Hum Brain Mapp 2016, 37:3337–3352.

72. Kalueff AV, Nutt DJ: Role of GABA in anxiety and depression. Depress Anxiety 2007, 24:495–517.

73. Nakamura S, Arima K, Haga S, Aizawa T, Motoi Y, Otsuka M, Ueki A, Ikeda K: Fibroblast growth factor (FGF)-9 immunoreactivity in senile plaques. Brain Res 1998, 814:222–225.

74. Aurbach EL, Inui EG, Turner CA, Hagenauer MH, Prater KE, Li JZ, Absher D, Shah N, Blandino P, Jr., Bunney WE, et al: Fibroblast growth factor 9 is a novel modulator of negative affect. Proc Natl Acad Sci U S A 2015, 112:11953–11958.

75. Lee KW, Yim HS, Shin J, Lee C, Lee JH, Jeong JY: FGF11 induced by hypoxia interacts with HIF-1alpha and enhances its stability. FEBS Lett 2017, 591:348–357.

76. Zhu T, Zhan L, Liang D, Hu J, Lu Z, Zhu X, Sun W, Liu L, Xu E: Hypoxia-inducible factor 1alpha mediates neuroprotection of hypoxic postconditioning against global cerebral ischemia. J Neuropathol Exp Neurol 2014, 73:975–986.

77. Shen Y, Gu J, Liu Z, Xu C, Qian S, Zhang X, Zhou B, Guan Q, Sun Y, Wang Y, Jin X: Inhibition of HIF-1alpha Reduced Blood Brain Barrier Damage by Regulating MMP-2 and VEGF During Acute Cerebral Ischemia. Front Cell Neurosci 2018, 12:288.

78. Li J, Wang Q, Wang H, Wu Y, Yin J, Chen J, Zheng Z, Jiang T, Xie L, Wu F, et al: Lentivirus Mediating FGF13 Enhances Axon Regeneration after Spinal Cord Injury by Stabilizing Microtubule and Improving Mitochondrial Function. J Neurotrauma 2018, 35:548–559.

79. Hu Y, Li L, Shen L, Gao H, Yu F, Yin W, Liu W: FGF-16 protects against adverse cardiac remodeling in the infarct diabetic heart. Am J Transl Res 2017, 9:1630–1640.

80. Miyake A, Chitose T, Kamei E, Murakami A, Nakayama Y, Konishi M, Itoh N: Fgf16 is required for specification of GABAergic neurons and oligodendrocytes in the zebrafish forebrain. PLoS One 2014, 9:e110836.

81. Boshoff EL, Fletcher EJR, Duty S: Fibroblast growth factor 20 is protective towards dopaminergic neurons in vivo in a paracrine manner. Neuropharmacology 2018, 137:156–163.

82. Amiri M, Braidy N, Aminzadeh M: Protective Effects of Fibroblast Growth Factor 21 Against Amyloid-Beta1-42-Induced Toxicity in SH-SY5Y Cells. Neurotox Res 2018, 34:574–583.

83. Chen S, Chen ST, Sun Y, Xu Z, Wang Y, Yao SY, Yao WB, Gao XD: Fibroblast growth factor 21 ameliorates neurodegeneration in rat and cellular models of Alzheimer’s disease. Redox Biol 2019, 22:101133.

84. Sun Y, Wang Y, Chen ST, Chen YJ, Shen J, Yao WB, Gao XD, Chen S: Modulation of the Astrocyte-Neuron Lactate Shuttle System contributes to Neuroprotective action of Fibroblast Growth Factor 21. Theranostics 2020, 10:8430–8445.

85. Ornitz DM, Itoh N: The Fibroblast Growth Factor signaling pathway. Wiley Interdiscip Rev Dev Biol 2015, 4:215–266.

