## Supplemental Data for "The FGF/FGFR system in the microglial neuroinflammation with *Borrelia burgdorferi*: intersectionality with other neurological conditions"

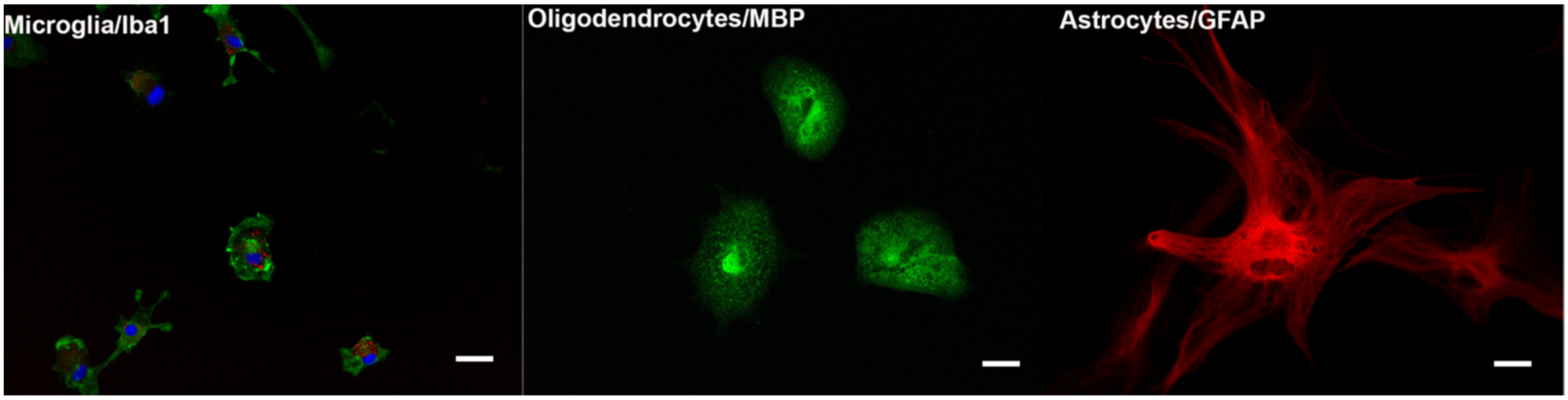


**SM1 Comparative sizes of primary rhesus microglia, primary rhesus oligodendrocytes and primary rhesus astrocytes in culture**

SM1 illustrates the size differences among the three main glial cells in the CNS**.** Primary rhesus glial cells were harvested and seeded as described in Methods. After microglial harvest and their seeding in 4 well chamber slides, the flasks containing cortex aggregate cultures were treated with 15-20mM Leucine methyl ester for 3h at room temperature, to remove any remaining microglia,. Cells were harvested with Trypsin EDTA treatment and seeded on 4-well chamber slides. Microglial cells were probed with both a mouse-Iba1 (1:10) and a rabbit Iba1 (1:100. Wako USA) antibodies. Mouse Iba1 was counterstained with secondary antibody conjugated with Alexa568 (red), while the rabbit antibody was captured with a secondary antibody conjugated to Alexa488 (green; both 1:1000). Oligodendrocytes were stained with a rabbit anti-human MBP (1:100) and a corresponding secondary antibody conjugated to Alexa488 (1:800). Astrocytes were stained with a mouse GFAP antibody conjugated to Cy3 (1:200). Microglial cells were stained by both antibodies, but the rabbit antibody stained more cells than the mouse antibody. However, all the cells stained by the mouse antibody were also stained by the rabbit antibody, indicating that the red staining cells were indeed Iba1 positive microglia. As seen in the immunofluorescent photographs, oligodendrocytes and astrocytes are much larger than microglia, when captured at the same magnification. Hence both size of the cells and Iba1 specificity were used to confirm microglia in cultures. Bar represents 25µm. Related to all the microscopy figures.

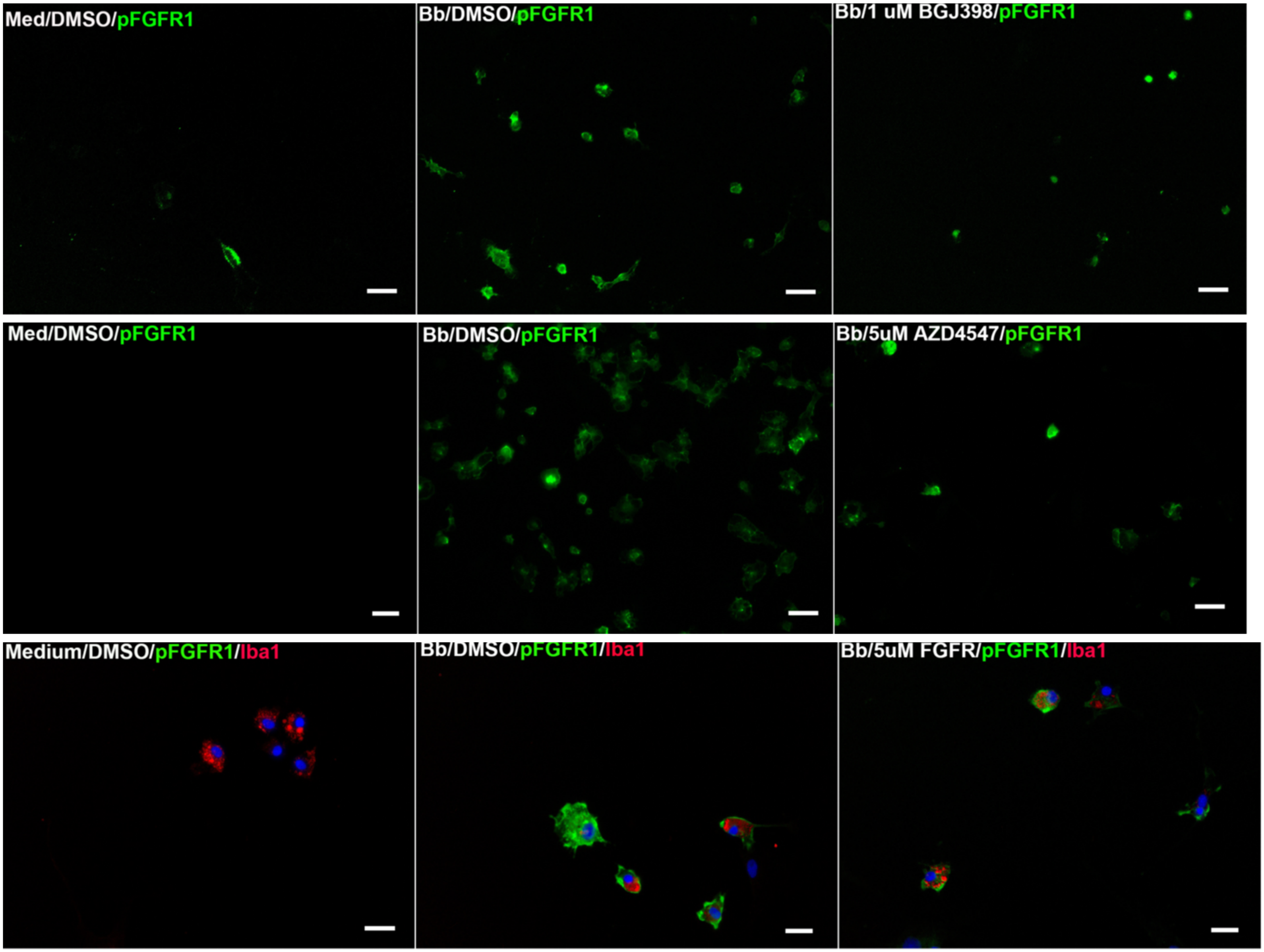


**SM2 Effectiveness of the inhibitors in downregulating FGFR1 signaling**

After treatment with respective inhibitors or DMSO for 24h, cells were fixed and stained for pFGFR1 (green) by immunohistochemistry. Treatment with triple FGFR inhibitors (BGJ398 and AZD4547) reduced both the intensity of staining as well as the number of cells staining positive for pFGFR1, while only the intensity was seen to be diminished in general with the FGFR1 inhibitor PD166866. Iba1 staining is in red while the nuclear stain is in blue. Bar represents 50µm for the top and middle panels, while for the bottom panel, bar is 25µm. Three separate experiments carried out on microglia derived from two different tissues is shown. Related to Figs. 2b and 3.

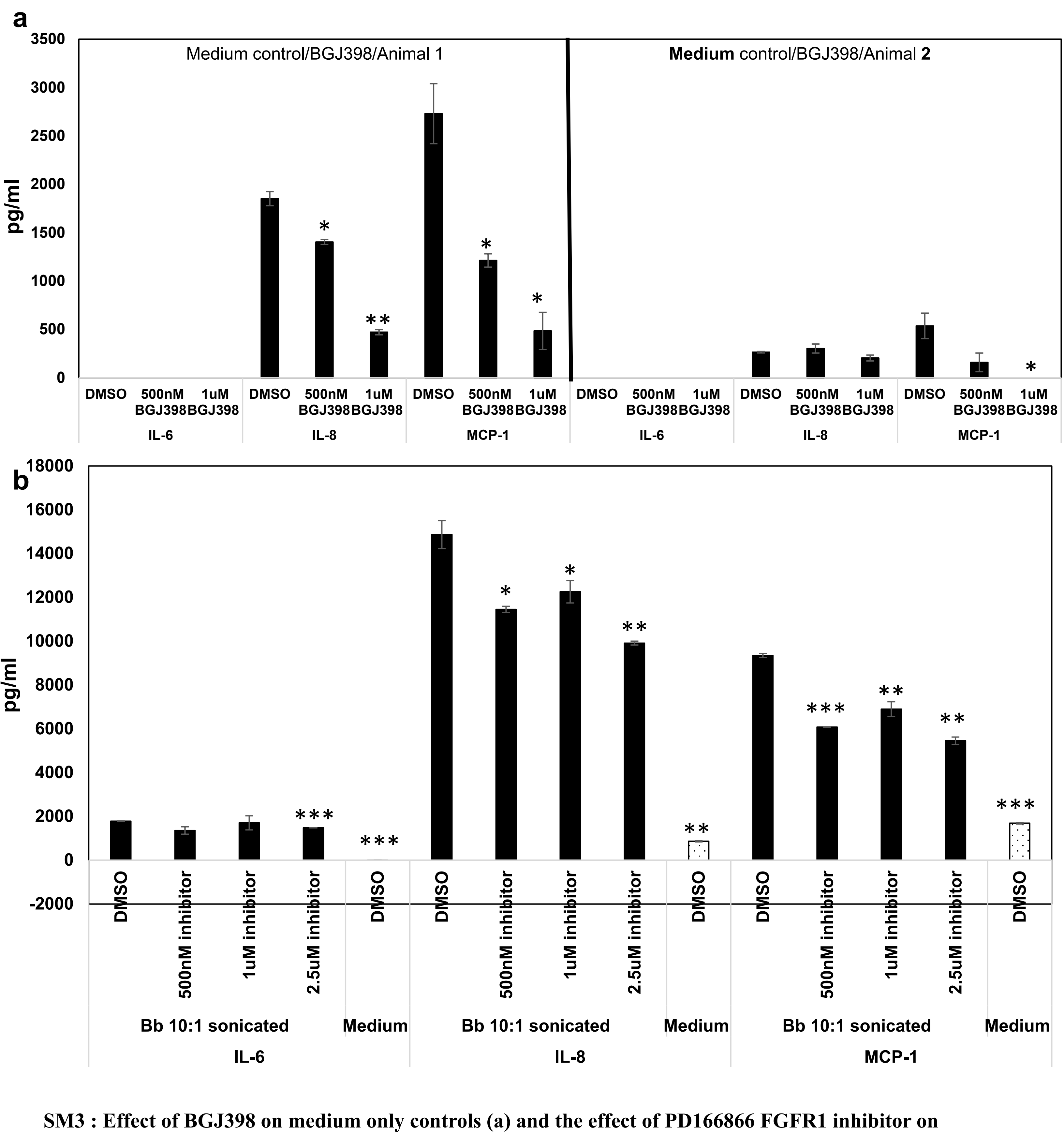


**SM3 : Effect of BGJ398 on medium only controls (a) and the effect of PD166866 FGFR1 inhibitor on inflammatory mediator output with non-viable *B. burgdorferi* (b)**

Primary rhesus microglia were pretreated with either DMSO/respective FGFR inhibitor for approximately 2hrs, prior to the addition of fresh Medium or sonicated bacteria. DMSO/inhibitor were left in with the Medium alone or sonicated remnants for another 24h, prior to supernatant collection and analysis by multiplex ELISA. Two experiments on microglia derived from two different brain tissues is shown in **a**., while a representative result of two experiments carried out on microglia derived from one brain tissue is shown in **b**. Bars represent standard deviation. Statistically significant results in comparison to Medium + DMSO (**a**) or Bb 10:1 sonicated + DMSO samples (**b**) are shown with black asterisks *$p < 0.05$, **$p < 0.01$, ***p<0001. Related to Table 1 and Fig.2b respectively.

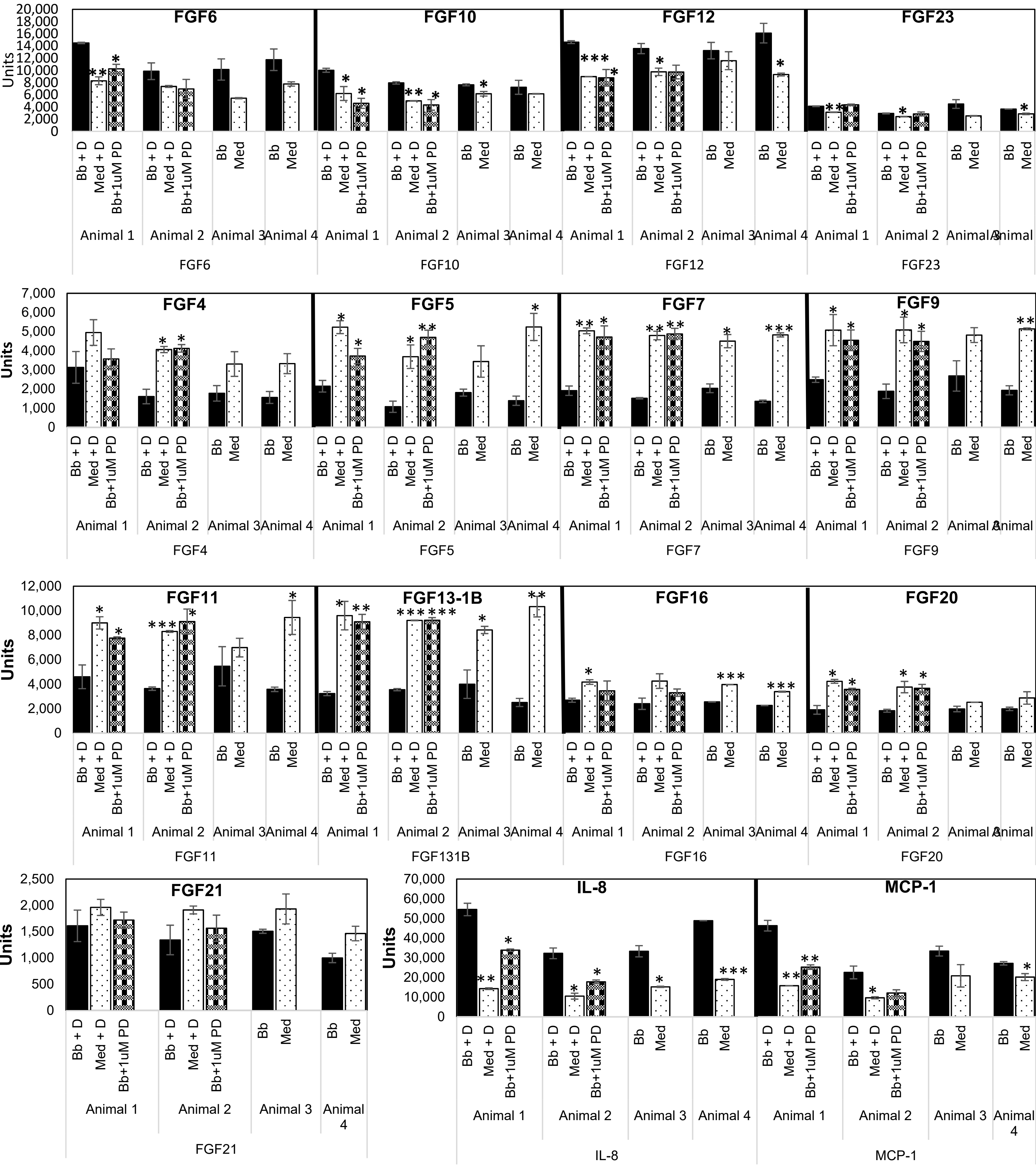


**SM4 Normalized data from the custom antibody array.** Data shows the salient upregulated and downregulated FGFs and the included positive controls IL-8 and MCP-1 . Animal 1, 2 3 and 4 represent Expts 1-4 in the heatmap. Bb-*B. burgdorferi* (10:1 MOI); M-Medium; D-DMSO; PD-PD166866. Bars represent standard deviation**.** Statistically significant results in comparison to Bb 10:1 or Bb 10:1 + DMSO samples are shown with black asterisks. *$p < 0.05$, **$p< 0.01$, ***$p<0001$. Related to Fig. 4b

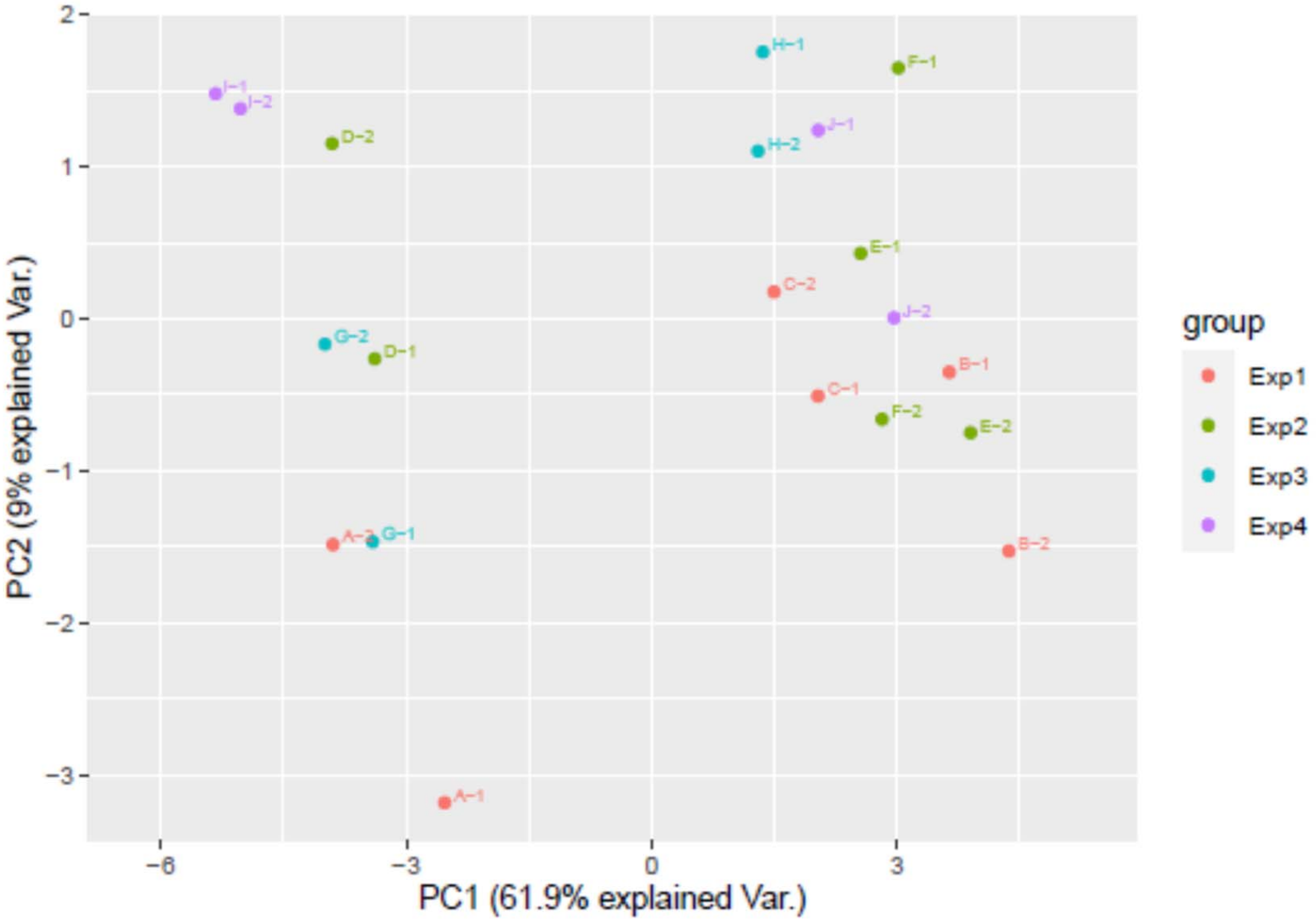


**SM5: Principal component analysis (PCA) plot of the 19 FGF biomarkers in 20 samples**

A, D,G,I are *B. burgdorferi* alone samples (with or without DMSO). B,E,H,J are Medium samples (with or without DMSO) and C,F are *B. burgdorferi* with 1µM FGFR1 inhibitor PD166866 samples.. PCA shows that *B. burgdorferi* group cluster away from Medium only group and *B. burgdorferi* with FGFR1 inhibitor group indicating distinct FGF responses between the 2 clusters. Related to Fig. 4b.

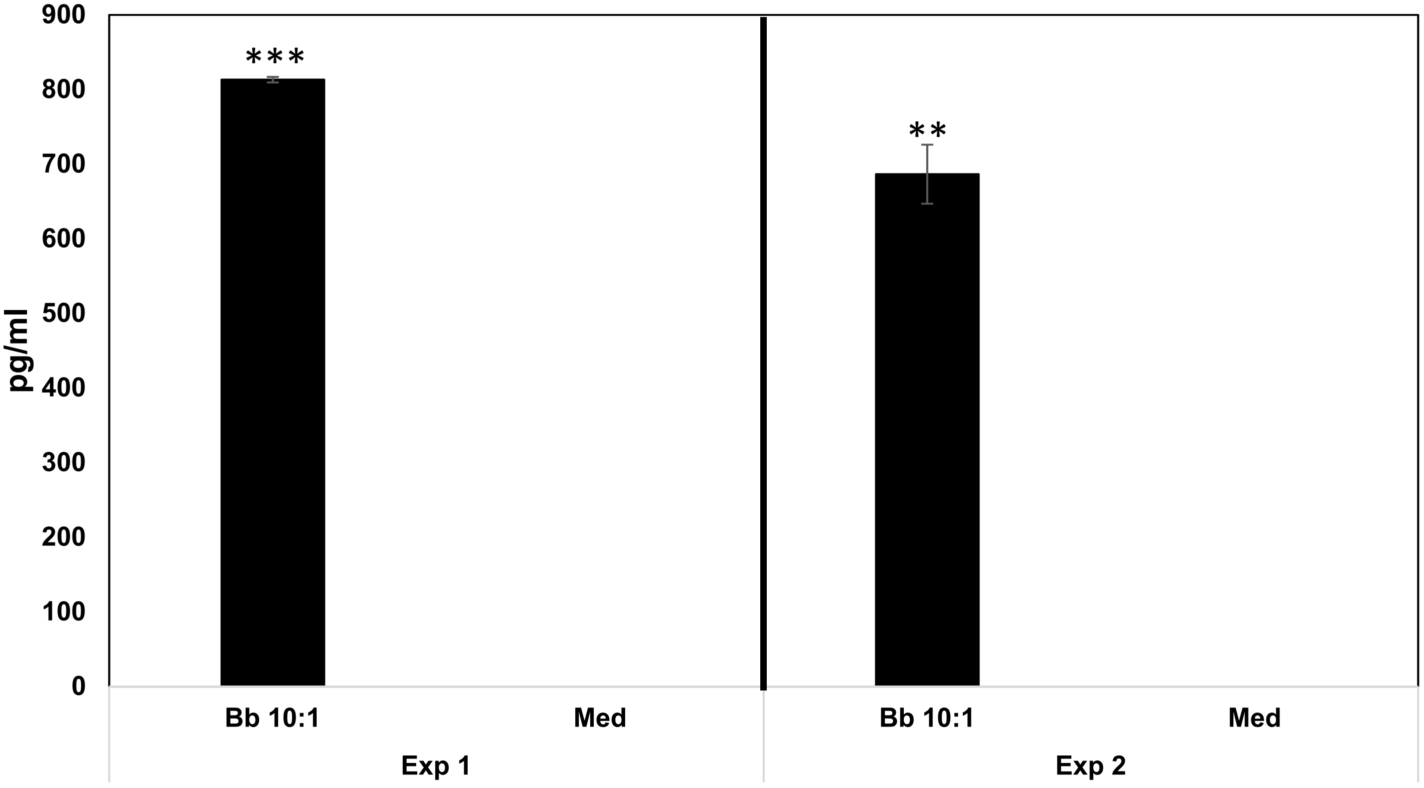


**SM6 Effect of *B. burgdorferi* exposure on FGF6 secretion from primary rhesus microglia**

Primary rhesus microglia were exposed to *B. burgdorferi* or medium controls for 24h. Supernatants were collected and analyzed for FGF6 by a human FGF6 ELISA. Two of the experiments conducted using microglia from one animal tissue are shown**.** Statistically significant results in comparison to Medium samples are shown with black asterisks. Bars represent standard deviation. *p < 0.05, **p< 0.01, ***p<0001. Related to Fig. 5a.
